# Assessment and Optimization of the Interpretability of Machine Learning Models Applied to Transcriptomic Data

**DOI:** 10.1101/2022.02.18.481077

**Authors:** Yongbing Zhao, Jinfeng Shao, Yan W Asmann

## Abstract

Explainable artificial intelligence aims to interpret how the machine learning models make decisions, and many model explainers have been developed in the computer vision field. However, the understandings of the applicability of these model explainers to biological data are still lacking. In this study, we comprehensively evaluated multiple explainers by interpreting pretrained models of predicting tissue types from transcriptomic data, and by identifying top contributing genes from each sample with the greatest impacts on model prediction. To improve the reproducibility and interpretability of results generated by model explainers, we proposed a series of optimization strategies for each explainer on two different model architectures of Multilayer Perceptron (MLP) and Convolutional Neural Network (CNN). We observed three groups of explainer and model architecture combinations with high reproducibility. Group II, which contains three model explainers on aggregated MLP models, identified top contributing genes in different tissues that exhibited tissue-specific manifestation and were potential cancer biomarkers. In summary, our work provides novel insights and guidance for exploring biological mechanisms using explainable machine learning models.

## Introduction

In recent years, many tools based on machine learning models have been developed and applied to biological studies, and most of which are developed for predictions. For example, AlphaFold was developed to predict protein 3D structure from amino acid sequences [1], P-NET was used to predict cancer treatment-resistance state from molecular data [2], and CEFCIG can predict cell identity regulators from histone markers [3]. Additionally, machine learning models can predict different biological features from the one single data type, depending which feature is paired with the input data when training the model. For instance, a variety of models have been developed to predict ncRNA [4], nucleosome [5], and chromatin accessibility/activity/states [6-9], all of which are from genome sequences.

Although these tools had been greatly successful in various biological topics, biologists are still curious about how a machine learning model makes decision, and which features of the input data play important roles in the model output. To answer these questions, explainable artificial intelligence (XAI) programs have recently emerged to enable the development of models that can be understood by humans [10, 11]. These XAI methods can also be applied to interpret machine learning models obtained from biological data by quantifying feature contributions to model prediction [12, 13]. The two most popular approaches to estimate the contribution of each input feature to the model output are: 1) Perturbing the input data and comparing outputs between the original and perturbated inputs; 2) Using backpropagation to measure the importance of each feature in the input data [14-16]. The former is intuitive but computationally expensive especially when exhaustively estimating all input features, and there is also the risk of underestimating feature contribution [17]. By contrast, the latter can measure the contribution of all input features in “one-shot”. Consequently many model explainers based on backpropagation were proposed and developed in the field of computer science and computer vision [18]. Benefiting from these model explainers, computational biologists discovered the syntax of transcription factor (TF) binding motifs by interpreting models trained to predict chromatin accessibility [19, 20]; and screened for cancer marker genes from models of cancer type classification [21-23]. There is no doubt that these explorations have showcased the potential of interpretable models in discovering meaningful biological mechanisms. However, a remaining problem is that results from different model explainers are highly variable [21]. Since these model explainers were not specifically designed for biological data, it is critical to evaluate their applicability in biology. Currently, there is still a lack of comprehensive understanding of these explainers in biological studies. To fill this gap, we optimized and assessed the performance of different model explainers and analyzed their biological relevance. To minimize the impact of model performance on the assessment of explainers, we tested explainers on well-trained models of predicting tissue type and cancer/normal status from gene expression data. In summary, this study provides a comprehensive guidance for optimization and applying interpretable machine learning to biological studies.

## Results

### Overview of model interpretability

In this study, we formulated a specific question to instantiate application of interpretable models to biological data. Can we quantify the contribution of individual genes to tissue type and disease status? Two steps were implemented to estimate attributes of each gene from the input sample. First, we built neural network models and trained the models with transcriptomes as input and sample tissue type and disease status as the prediction output. Models were built based on two types of neural networks, convolutional neural network (CNN) and multiple layer perceptron (MLP) (model architectures are detailed in the Methods section). In general, CNN is more complex than MLP. Next, we applied model explainers to compute quantitative score of each gene’s contribution to the model’s prediction, which are named gene contribution scores. We tested eight popular model explainers and their vriations commonly used in computer vision, and then assessed and compared their applicability and performances on each pretrained model. These explainers are: Saliency, InputXGradient, GuidedBackprop, IntegratedGradients, DeepLift, DeepLiftShap, GuidedGradCam, and GuidedGradCam++ (**Table S1**) [17, 18, 24-29]. Since GuidedGradCam, and GuidedGradCam++ were developed for CNN specifically, only the first six explainers were tested on MLP.

We used 27,417 RNA-Seq samples from GTEx and TCGA projects to train CNN or MLP-based models. These samples were isolated from 82 distinct normal and cancer tissues and cell types (**Table S2**). After training, the prediction accuracy of all models is comparable, with a median value of 97.2% for CNN and 97.8% for MLP. The convolutional layers of the CNN models require that, as input data, gene expression values should be organized with a fixed gene order in a 2-D matrix. Therefore, we tested various gene orders for the CNN-based models, e.g., sorting genes according to their genomic coordinates [22]. Results indicated that gene order did not affect model performance in terms of prediction accuracy. Although models were trained with all 82 different normal and cancer tissues, results from four normal tissues (liver, lung, ovary and pancreas) are reported here to illustrate the applicability and performance of the eight explainers.

### Direct use of explainers from computer vision resulted in poor reproducibility

Randomness is often challenging in machine learning, which is present in both model training and model interpretation [30, 31]. For this reason, we measured both intra-model and inter-model reproducibility of each explainer. During the testing, each explainer was applied to pretrained models based on CNN or MLP respectively.

First, we tested intra-model reproducibility by applying an explainer to the same pretrained model 5 times (5 replicates per model per explainer) and for each explainer we checked correlations of gene contribution scores as well as pair-wise overlap of the top 100 contributing genes between replicates. We found that intra-model reproducibility, in terms of Spearman’s correlation of gene contribution scores and overlap of gene IDs among the top 100 contributing genes, are low for most explainers on both CNN and MLP-based models (results from normal liver samples: **Figure 1**, results from normal lung, ovary, and pancreas: **Figure S1**). The exception was GuidedGradCam++, which was previously used to identify cancer markers [21].

**Figure 1.**
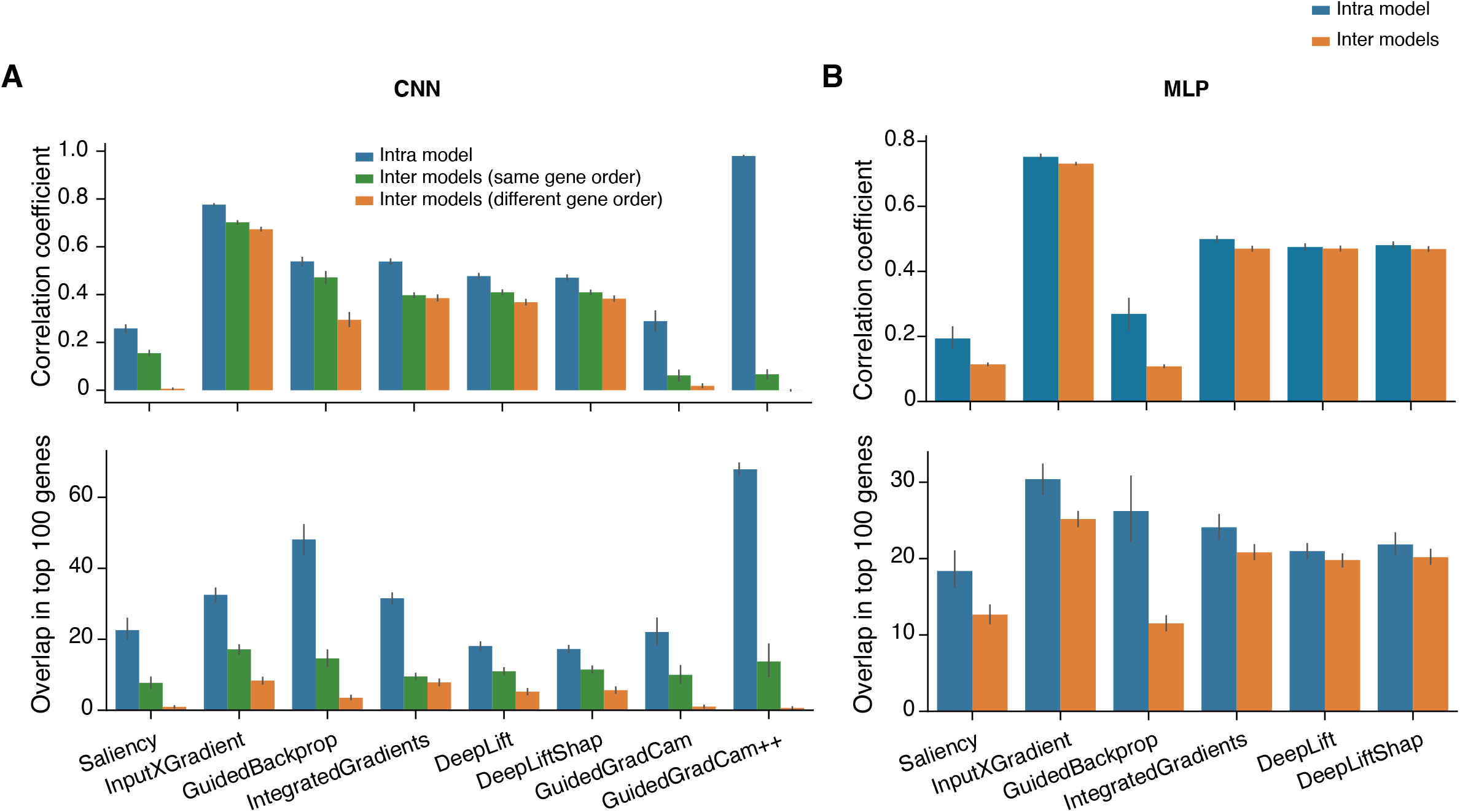
Performance of different model explainers without optimization. **A**. Spearman’s correlation of gene contribution scores (upper panel) and overlap in the top 100 contributing genes (lower panel) in liver among replicates from the same pretrained model, different pretrained models with the same gene order, and different pretrained models with different gene orders on CNN-based models. **B**. Spearman’s correlation on gene contribution scores (upper panel) and overlap in the top 100 contributing genes (lower panel) in liver among replicates from the same pretrained model and different pretrained models on MLP-based models. CNN: Convolutional Neural Network, MLP: Multilayer Perceptron.

As shown in **Figure 1**, we also tested inter-model reproducibility by applying each explainer to five different models with comparable prediction accuracies (5 models per explainer). These five models were trained using the same model architecture and training data set but with slightly different hyperparameters. For CNN models, we also tested the impact of different input gene orders since CNN requires organizing input genes in a 2D matrix. The inter-model reproducibility of all explainers, including GuidedGradCam++, were significantly lower than those from the intra-models. For CNN-based models, even though the gene order had little impact on prediction accuracy, it had significant impact on the model reproducibility especially regarding the overlap of top contributing genes. For MLP-based models, the reproducibility of intra-model and inter-model tests were similar, however both Spearman’s correlation and overlap of the top 100 contributing genes were very low.

In summary, gene contribution scores vary greatly intra- and inter-models for both CNN and MLP. These tests were performed per explainer. We expect that reproducibility across different explainers would be much worse. Therefore, model explainers developed for computer vision may not be directly applied to answering biological questions. Model interpretability in computer vision aims to identify visual features consisting of multiple similar pixels in an area and variations within the area have limited impact on outcome. However, interpreting biological data such as the transcriptomes requires single gene resolution since genes within an area were arbitrarily placed together, therefore the results are very sensitive to random noise.

### Optimization on model interpretability

Since it is not feasible to directly transfer model explainers from computer vision to biology, we tested whether these explainers can be optimized and adjusted for biological data. First, we borrowed a de-noising strategy widely used in computer vison, “SmoothGrad: removing noise by adding noise” [32]. Instead of estimating gene contribution scores in a sample in “one pass”, SmoothGrad calculates gene contribution by averaging contribution scores from multiple explanation estimates per sample by adding random noise into the expression data each pass. Unfortunately, the strategy of SmoothGrad did not improve inter-model reproducibility. On the contrary, it lowered the performance of all explainers on both CNN and MLP except for Saliency on MLP and GuidedBackprop on CNN (**Figure 2A and Figure S2)**. For Saliency on MLP, the improvement saturates when the number of repeat estimates reached 50, while performance of GuidedBackprop plateaued after 30 repeats. Next, we tested whether repeating explanation without adding random noise into the expression data, which we defined as simple repeat, would be beneficial. Results of the simulation indicated that repeats without adding noise significantly improved the performance of all explainers on both CNN and MLP, except for Saliency and GuidedGradCam++ on CNN (**Figure 2B and Figure S3**). For most explainers, improvement saturated after 20 repeats for CNN, and after 40 repeats for MLP.

**Figure 2.**
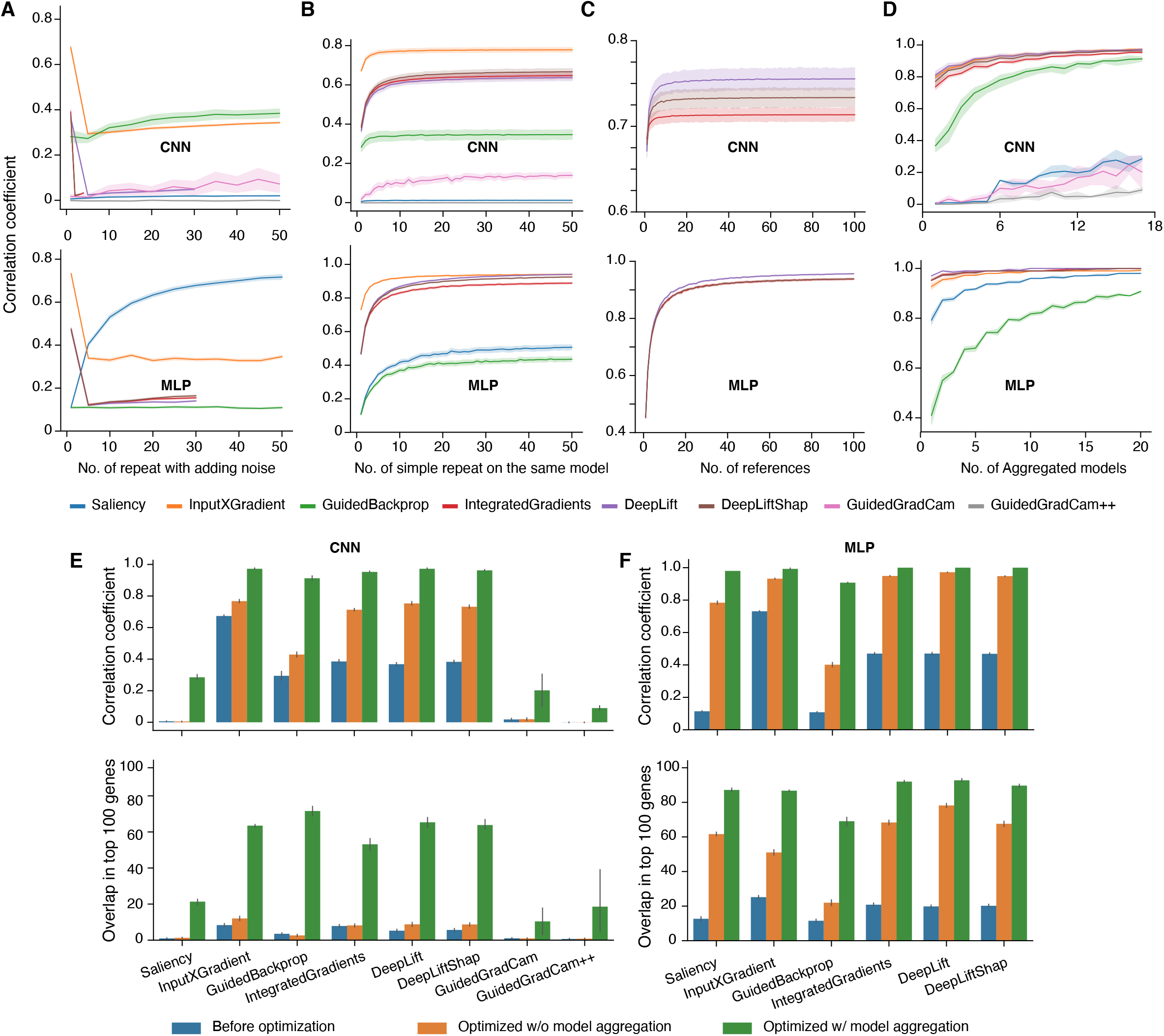
Optimization of different model explainers. Spearman’s correlation of gene contribution scores in liver from CNN models (upper panels) and MLP (lower panels). **A**. Performance by averaging contribution scores from multiple estimates per sample by adding random noise into the expression data each pass; **B**. Performance from running the same model multiple times and averaging contribution scores without adding random noise (simple repeats); **C**. performance of repeated reference zero for CNN, and number of references randomly selected from 2000 simulated reference universal for MLP models, for three explainers that require reference samples; **D**. Performance of model aggregations. **E**. Spearman’s correlation on gene contribution scores (upper panel) and overlap in the top 100 contributing genes (lower panel) among replicates from different pretrained models with different gene orders based on CNN-based models. The analyses were carried out with three different optimization strategies: without optimization, with optimized conditions for each explainer but without model aggregation, and with optimized conditions for each explainer and with model aggregation. **F**. same as E) but based on MLP-based models.

DeepLift, DeepLiftShap and IntegratedGradients require a reference baseline when estimating gene contribution scores, where a reference is a synthetic, randomly generated transcriptome. In computer vision, a black image (all-zeros input) or random pixel values are often used as references, and for motif identification of regulatory elements, scrambled genomic sequence is demonstrated as good references [17]. In this study, we compared four types of references, named as reference zero, normal, universal, and specific. Reference zero and normal are equivalent to black image and random pixel values respectively. For reference universal and specific, we estimated mean (µ) and standard deviation (σ) of each gene’s expression level across samples, and then randomly generated a value based a truncated normal distribution N (µ, σ). Reference universal uses samples from all 82 tissue types while reference specific only uses samples from a specific tissue type.

We tested performance of these four kinds of references individually, as well as in combination to evaluate whether multiple references would improve reproducibility among models. First, for individual references, simulation results showed that reference zero is the best for CNN-based models, while universal is preferred on MLP-based models (**Figure S4)**. The result is consistent on all the three explainers that required references. Next, we tested the impact of multiple references on reproducibility. Since effect of using multiple references with zero is equivalent to that of simple repeat with single reference zero, these two kinds of optimization, using multiple references zero and simple repeat, cannot contribute to reproducibility additively. Therefore, we compared reproducibility by combing simple repeat with multiple references as reference normal, universal, or specific, with the reproducibility by combing simple repeat with single reference zero (which is equivalent to multiple references zero without simple repeat). Interestingly, we found that reference zero still outperformed the other three kinds of references on CNN-based models (**Figure S5)**. Similar as simple repeat, improvement saturates when number of references zero reach to 20 on CNN-based models, while the number of references universal goes to 60 on MLP-based models (**Figure 2C**).

Since that the intra-model reproducibility was significantly improved by repeating the explanation process multiple times and averaging contribution scores from different estimates (**Figure 2B**), we next tested the benefits of inter-model aggregation. Therefore, we applied optimized parameters of each explainer on CNN or MLP-based models (**Table S2**), estimated gene contribution scores on each pretrained model individually, and then averaged inter-model results. Indeed, aggregating models significantly increased reproducibility (**Figure 2D, and Figure S6**). Especially, Spearman’s correlations for DeepLift, DeepLiftShap, IntegratedGradients, and Saliency reached nearly 1.0 on MLP. In general, the reproducibility of all explainers was significantly increased on both CNN and MLP-based models after aggregating models (**Figure 2D-F, and Figure S7**). Of note, model aggregation has extremely strong impact and improvement on the reproducibility of all explainers on CNN-based models in terms of overlaps between the top 100 contributing genes (**Figure 2E**, lower panel). For most explainers on MLP-based models, Spearman’s correlations on gene contribution scores based on model aggregation were higher than 0.9, and over 90% of top 100 contributing genes overlapped between replicates on the same explainer (**Figure 2F**).

To summarize, gene contribution scores were highly reproducible from the same explainer with optimized parameters. Reproducibility of the top 100 contributing genes was better on MLP-based models than those on CNN-based models. One possible reason is that CNN-based models are much more complex than MLP-based models and can be hard to be interpreted.

### Consistency across model explainers

So far, we’ve tested the performance within each explainer. To test consistency of gene contribution scores across different explainers, we checked the overlap of the top 100 contributing genes identified by different explainers with and without model aggregation (**Table S3**). Within CNN or MLP models, model aggregation did not only improve reproducibility within the same explainer, but also across explainers. However, the top 100 contributing genes from CNN-based models with model aggregation did not overlap with those from MLP-based models with or without model aggregation. Moreover, within CNN-based models, the top 100 contributing genes with model aggregation did not overlap with those without model aggregation either, which suggests that model aggregation resulted in explainers identifying completely different set of genes. By contrast, top contributing genes from MLP-based models were highly consistent with and without model aggregations. We further explored why model aggregation had different impact on MLP and CNN. We first defined the top 100 contributing genes from optimized parameters with model aggregations as the baseline of comparison for each explainer. Next, we compared the top 100 contributing genes from each explainer without optimization to the baseline. It was found that, without optimization on MLP-based models, top contributing genes shared by two or more replicates had higher overlap with the baseline than that between top contributing genes from individual replicates and the baseline (**Figure S8**). However, this conclusion was not seen on CNN-based models. This result suggests that the top contributing genes from different MLP models are convergent, while those from different CNN models are very divergent.

Intriguingly, the measurement of reproducibility highlighted three representative groups each with different explainers, model types (CNN or MLP), and aggregation status (**Figure 3**). The three groups are group I: DeepLift, DeepLiftShap, GuidedBackprop, InputXGradient, and IntegratedGradients on CNN-based models with model aggregation; group II: DeepLift, DeepLiftShap, InputXGradient, and IntegratedGradients on MLP-based models with model aggregation; and group III: GuidedBackprop and Saliency on MLP-based models with model aggregation. Next, we delved into the top contributing genes identified by these three groups.

**Figure 3.**
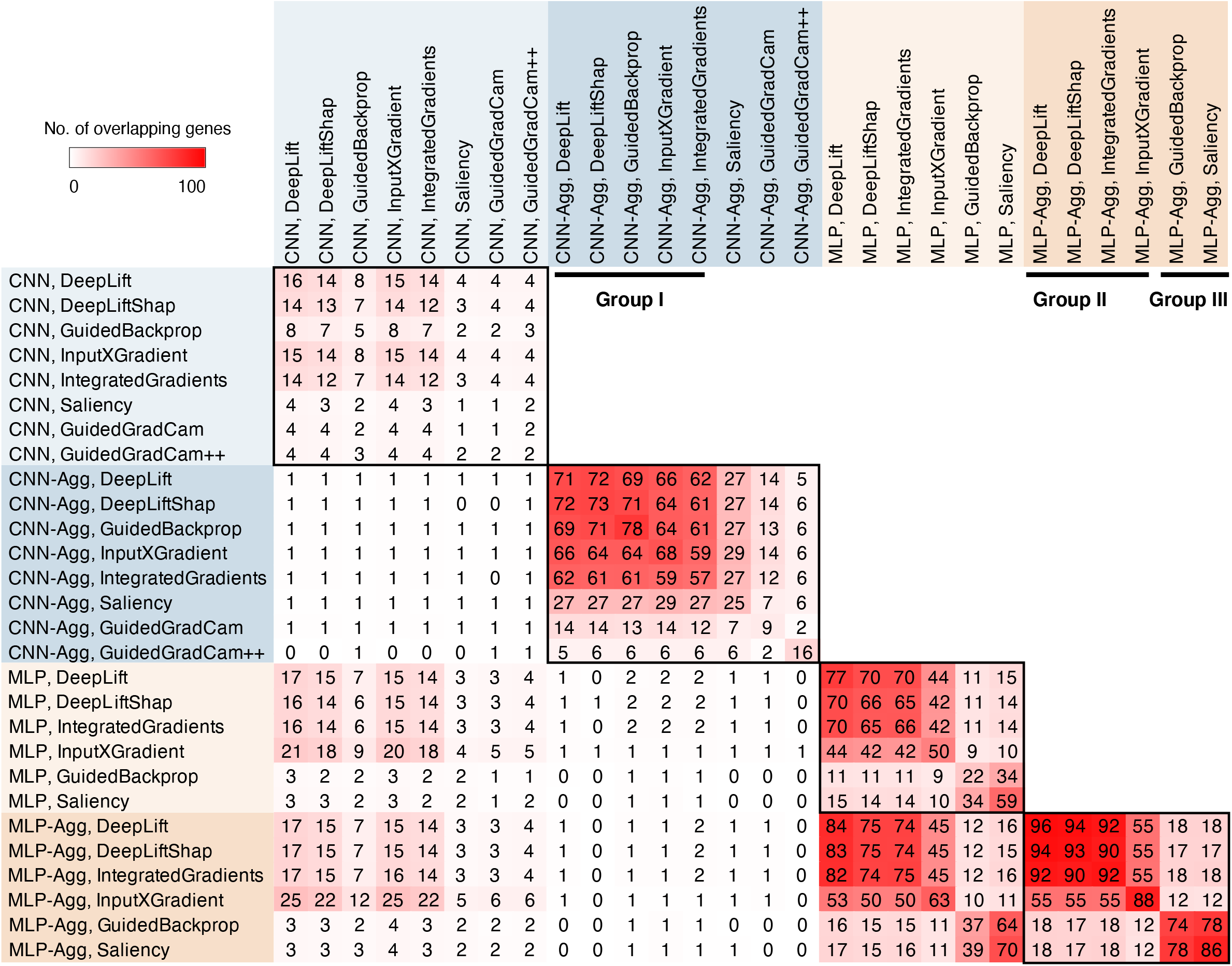
Overlap of the top 100 contributing genes across explainers with and without model aggregation. Three representative groups (I, II and III) are marked by black bars. Aggregation of CNN models or MLP models are abbreviated as CNN-Agg or MLP-Agg.

### Expression status of top contributing genes

It is important to understand the biological relevance of the top contributing genes by different model architecture (MLP vs. CNN) and different explainers. Since contribution scores were derived from the input gene expression values, we first calculated Spearman’s correlation between gene contribution scores and expression levels (**Figure S9**). We expected high correlation for genes identified by InputXGradient because gene expression level is a cofactor in computing gene contribution scores by InputXGradient. Indeed, we found weak correlations of all explainers in both group II and group III except for InputXGradient. Conversely, strong correlations were observed from 4 explainers in group I: InputXGradient, IntegartedGradients, DeepLift, and DeepLiftShap. However, it is puzzling that GuidedBackprop from Group I showed negative correlations for unknown reasons.

Additionally, we checked overlaps between the top 100 contributing genes and the top 100 expressed genes for all explainers on both CNN and MLP-based models (**Figure S10**). In liver, nearly 50% of the top contributing genes from group II overlapped with top expressed genes, while the overlaps were less than 10% in groups I and III (**Figure 4A**). Group II, as described above, are DeepLift, DeepLiftShap, InputXGradient, and IntegratedGradients on MLP-based models with model aggregation. Strikingly, model aggregation eliminated the already moderate overlaps in CNN-based models (Group I). Another noticeable finding for group I is that though Spearman’s correlation between gene contribution scores and expression level were very high, majority of the top contributing genes were not highly expressed.

**Figure 4.**
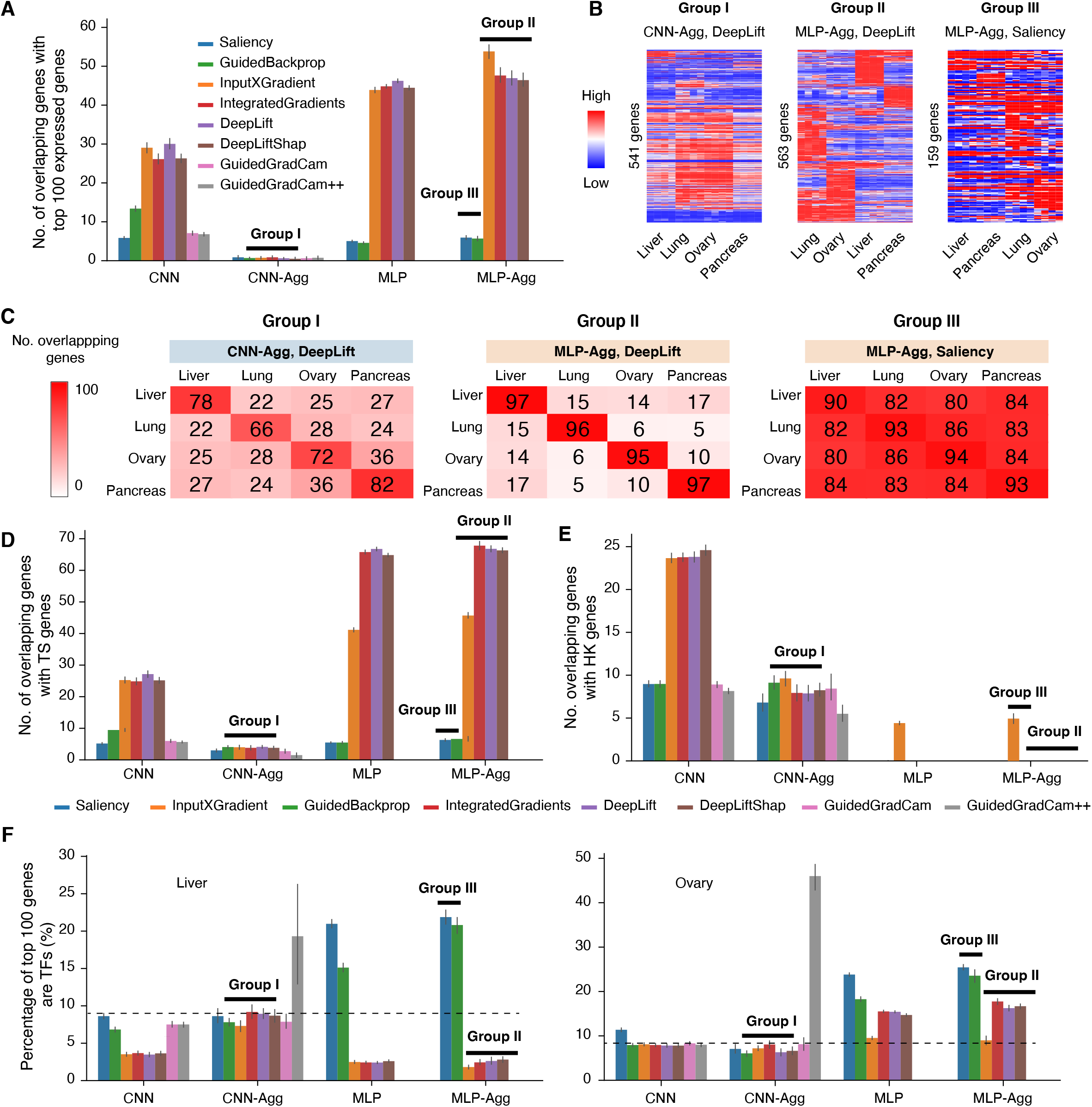
Biological relevance of the top 100 contributing genes. **A**. Overlaps between the top 100 contributing genes and the top 100 highest expressed genes in liver samples. Explainers from group I, II and III are marked by with black bars. **B**. Expression profiles of the top 100 contributing genes in liver, lung, ovary and pancreas identified by DeepLift on CNN-based models with model aggregation (representative of Group I), DeepLift on MLP-based models with model aggregation (Group II), and Saliency on MLP-based models with model aggregation (Group III). **C**. Overlaps in the top 100 contributing genes among liver, lung, ovary and pancreas identified by DeepLift on CNN-based models with model aggregation (Group I), DeepLift on MLP-based models with model aggregation (Group II), and Saliency on MLP-based models with model aggregation (Group III). **D**. Overlaps between the top 100 contributing genes and tissue-specifically (TS) expressed genes in liver samples. **E**. Overlaps between the top 100 contributing genes and housekeeping (HK) genes in liver samples. **F**. Percentage of the top 100 contributing genes are transcription factors (TFs) in liver samples (left panel) and ovary samples (right panel). Dashed lines indicate the overlap percentage by random chance.

Since different tissues have distinct phenotypes, we wondered whether the top contributing genes of different tissue types exhibit distinct expression profiles. Heatmap of the top contributing genes clearly demonstrated tissue-specific manifestations for group II (represented by DeepLift on MLP with model aggregation), and the patterns were much weaker in both group I (represented by DeepLift on CNN with model aggregation) and group III (represented by Saliency on MLP with model aggregation) (**Figure 4B**). In addition, the total number of genes from group III is much lower than that of both group I and II, after removing redundant genes from the top 100 contributing genes across tissues. This suggests that the top 100 contributing genes were largely shared across tissues in group III, which was validated by comparison across tissue types in all explainers (**Table S4**). Among the three groups, the top contributing genes in both groups I and II are tissue specific, while the top contributing genes in group III are highly shared across tissues (**Figure 4C**).

Considering that the top contributing genes in group I and II were mostly tissue specific, we are curious how the top contributing genes are related to tissue-specifically (TS) expressed genes. For this purpose, we identified TS genes across all 82 tissues and cell types, which were used in model training. It was found that about 70% of the top contributing genes overlap with TS genes in group II in liver (**Figure 4D**). The fractions vary across tissues (**Figure S11A**), since there are different numbers of TS genes in each tissue type (**Figure S11B**). The percentages drop to less than 10% in both group I and group III. Interestingly, model aggregation also diminishes overlaps with TS genes in most explainers on CNN-based models.

In addition, we observed that many of the top contributing genes (in group I particularly) are expressed at comparable levels across tissues, we investigated the relationships between the top contributing genes and housekeeping (HK) genes. Results showed that about 10-20% of top contributing genes overlap with HK genes in group I, and the overlap was also further reduced by model aggregation (**Figure 4E, and Figure S12**). Conversely, no overlap was found from both group II and group III, except for the explainer InputXGradient.

### Enrichment of top contributing genes in biological functions

To understand the biological functions of the top contributing genes, we performed Gene Ontology (GO) enrichment analysis. No enrichment was found on genes identified by all explainers in group I. The enriched GO terms by genes from group II were mostly unique for each tissue type and tissue-specific functions (**Table S5**). For example, enriched GO terms in liver are molecular function related to lipoprotein and lipoprotein lipase activities, while GO terms enriched in pancreas are associated with binding of oligosaccharide, peptidoglycan and so on. Additionally, in group II, results between DeepLift, DeepLiftShap and IntegratedGradients are slightly more agreeable compared to that from InputXGradient. For group III, we expected similar GO terms enriched across tissue types since top contributing genes from different tissues highly overlapped. This turned out to be the case. GO enrichment analysis showed that top contributing genes in group III are enriched in CCR7 chemokine receptor binding, neuropeptide hormone activity, neuropeptide receptor binding, and DNA-binding transcription activator activity across tissues. Next, we checked how top contributing genes are related to transcription factors (TFs) and TF cofactors. We found about 20 genes overlapping with TFs in group III, which is more than 2-fold enrichment than by random chance (**Figure 4F**). By contrast, genes in group II showed depletion of TFs in liver, but 1.5-fold enrichment in Ovary (**Figure 4F, and Figure S13**). No enrichment or depletion was found in group I, except for genes identified by GuidedGradCam++. As for TF cofactors, there are low overlap in all three groups (**Figure S14**).

### Top contributing genes in cancers

Group II’s top contributing genes are tissue specific with tissue-specific manifestations of expression values. Therefore, we asked how the expression pattern of the top contributing genes changed from normal to cancer tissues. We compared normal and cancer samples of liver, lung, ovary and pancreas from GTEx and TCGA, and studied the expression differences of top contributing genes. About 40-80% of the top contributing genes were differentially expressed genes between normal and cancer tissues identified by DeepLift on MLP (group II) which is about twice more than the random chance (**Figure 5A**). The percentages ranged from 30% to 60% by Saliency on MLP (group III) which is about 1.5-fold over random chance. Group I was not included in this analysis since model aggregation eliminated many features in common from different explainers on CNN-based models and no biological enrichments were found in the top contributing genes.

**Figure 5.**
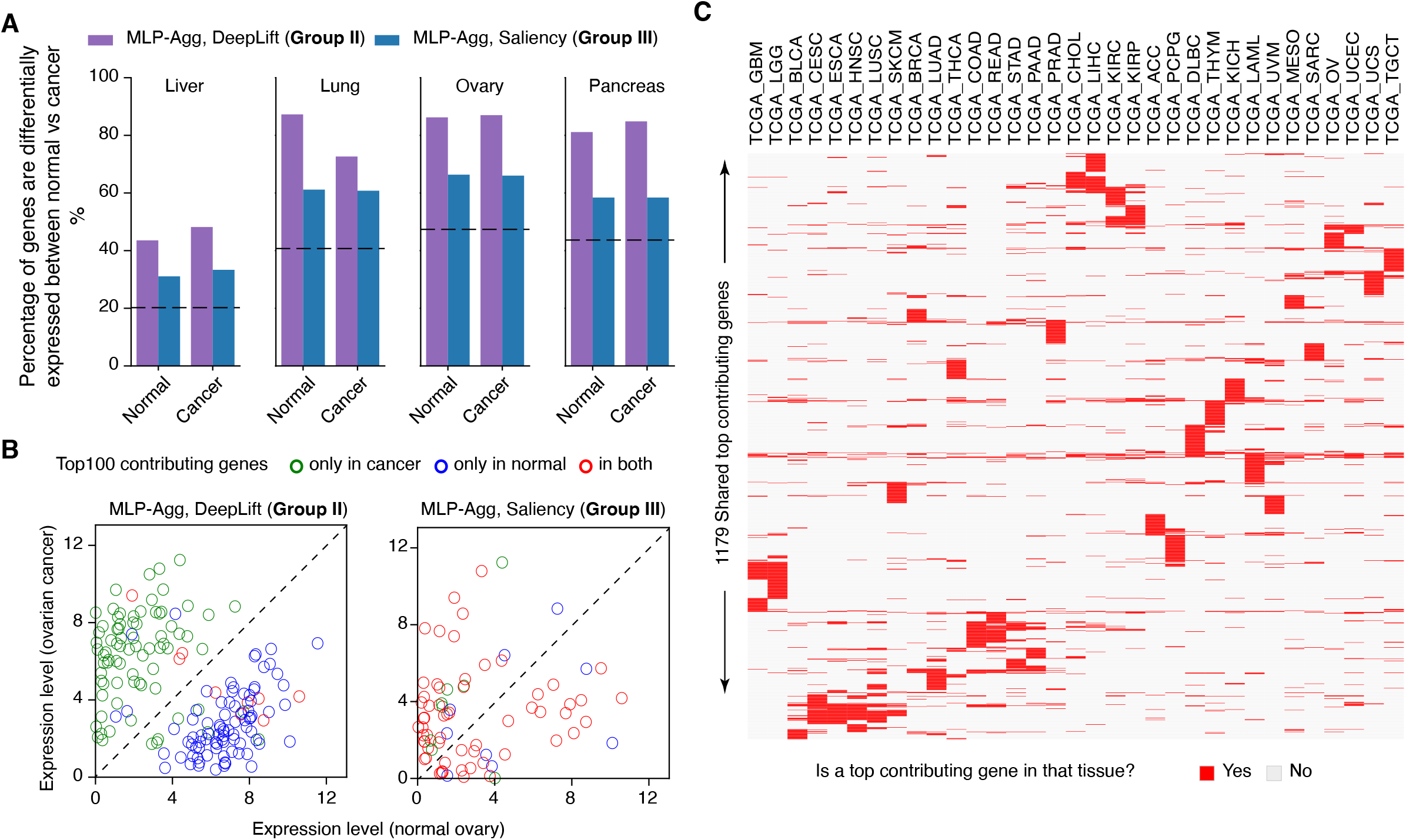
Top contributing genes in cancers. **A**. Percentages of the top 100 contributing genes are differentially expressed between normal and cancer tissues. Dashed lines indicate the percentage of differentially expressed genes by random chance. **B**. Expression levels of the top 100 contributing genes observed only in cancer samples, only in normal samples, and in both normal and cancer samples, of genes identified by DeepLift on aggregated MLP models (MLP-Agg, representative of Group II, left panel) and by Saliency on aggregated MLP models (representative of Group III, right panel). **C**. Heatmap of the 1179 shared top contributing genes from 33 TCGA cancer types demonstrated tissue specificity.

Interestingly, differentially expressed top contributing genes are segregated into two distinct populations in group II (**Figure 5B, and Figure S15**). Specifically, the top contributing genes specific to normal tissues are downregulated in cancer, while those specific to cancer are upregulated in cancer. For example, Glypican-3 (*GPC3*), a member of heparan sulfate proteoglycans family, is one of top contributing genes in liver cancer but not in normal liver. *GPC3* is often observed to be highly elevated in hepatocellular carcinoma and is a target for diagnosis and treatment of hepatocellular carcinoma [33]. However, similar segregation was not found in group III (Saliency on MLP-based models) because top contributing genes from group III were mostly shared between normal and cancer tissues. Together, the expression profiles suggest that the top contributing genes in group II might be potential cancer biomarkers. We identified the top 100 contributing genes in individual samples of all 33 different cancer types and named those shared by two or more samples of the same cancer type as shared top contributing genes. In total, 1179 genes were identified as shared top contributing genes. As expected, these shared top contributing genes are mostly tissue-specific (**Figure 5C**). Among these shared top contributing genes, we further studied the known oncogenes and tumor suppressor genes based on annotation in OncoKB [34]. Heatmap analysis showed that some oncogenes and tumor suppressor genes are shared by multiple cancer types, such as *SFRP2*, while the others are specific to one or very few cancers (**Figure S16**).

## Conclusions

The beauty of interpreting machine learning models is that it converts the complex mathematical rules learned by neural networks into biological rules and provides new insights into biology. To facilitate application of interpretable machine learning model, we established a series of optimization steps and compared the biological relevance of different model explainers. Since the tests in this study were based on models that predict tissue types from transcriptomes, applications using other types of biological data may require further investigations. In addition, even though the current optimizations demonstrated good performance on MLP-based models for a subset of explainers, some important genes may still be missing from the top contributing genes. For example, a machine learning model might choose only one of two highly correlated genes to use for prediction. Alternatively, the contribution of two highly correlated genes might be diluted if the model chooses to use both genes and thus, the contribution here might not reflect actual biological importance. These factors might partly explain the low reproducibility of individual single models and why improvement could be made by aggregation of models. Overall, we believe this study will provide novel insights to optimize interpretable machine learning in biological studies.

A recent paper pointed out five potential pitfalls of applying machine learning in genomics: 1) distributional differences; 2) dependent examples; 3) confounding; 4) leaky pre-processing; and 5) unbalanced class[35]. These technical challenges of applying machine learning models to genomics data are nontrivial and should be paid close attention to in addition to the optimization strategies we laid out in this study.

Typically, complicated models are not easily interpretable [36], which is also confirmed by the poor performance when interpreting CNN-based models. In this study, the optimized strategy significantly increased the interpretability on MLP-based models for a subset of explainers, but not on CNN-based models for any explainers. The aggregated CNN model approach should perhaps be categorized into a new modeling strategy, which is similar to “averaging” of models. The “averaging” strategy indeed mitigated randomness to some extent but didn’t show biological relevance. Therefore, even if models of different architectures had comparable prediction performances, it’s probably preferable to use models with relatively simpler architectures for model interpretation.

The top contributing genes detected by explainers in group II (DeepLift, DeepLiftShap, InputXGradient, and IntegratedGradients on MLP-based models with model aggregation) exhibited tissue-specific manifestation in both gene ontology and expression profile, which is expected based on prior knowledges about tissue specificity and cell identity [37-40]. Therefore, explainers in group II are more suitable for biological study, especially when exploring biological questions based on transcriptomic data. In recent years, single-cell RNA-Seq technology has been widely applied to different tissue and diseases, leading to the discovery of many well-defined sub-cell populations [41, 42]. Although this study assessed model interpretability on bulk RNA-Seq transcriptomes, the optimization strategies proposed here can also be applied to single-cell transcriptomes to quantify individual gene contribution and identify important genes in each sub-population. It is expected that interpretable machine learning models will also benefit understandings on tissue heterogeneity, disease mechanisms and cellular engineering at single cell resolution.

## Methods

### Human transcriptome collection and processing

Total of 27,417 RNA-Seq samples were used in our study, among which 17,329 and 10,088 RNA-Seq samples were collected from GTEx and TCGA, respectively [43]. These samples are from 47 distinct primary normal tissues and 2 cell lines (with prefix GTEx_ in the tissue code) and 33 primary tumors (with prefix TCGA_ in the tissue code). Pre-processed TCGA and GTEx RNA-Seq gene expression level data were downloaded from GTEx Portal (phs000424.v8.p2) and Recount2 database [44] respectively. For TCGA data, only primary tumor samples were included. Tissue types (normal or cancer) names remained the same as defined by TCGA and GTEx Projects. For each sample, the expression levels of 19,241 protein-coding genes were normalized to log2(TPM + 1) and then used for analyses.

### Architecture of CNN models

We used a five-layer convolutional neural network to build the CNN models, which included three convolutional layers, one global average pooling layer and one fully connected layer sequentially. Each layer included 64, 128, 256, 256 and 82 channels respectively. The kernel sizes for the three convolutional layers were 5, 5, and 3 respectively, and each convolutional layer is followed by max pooling with kernel size of 2. Batch normalization and rectified linear unit activation function (ReLU, which can be presented as f(x) = max (0, x)) were applied immediately after max pooling of each convolutional layer and global average pooling layer.

As the input of the CNN model, normalized expression values of 19,241 protein-coding genes from a sample were transformed into a 144×144 matrix, and zero-padding was used at the bottom of the matrix. The final fully connected layer produced an vector of 82-probability-like scores, each corresponding to one of the 82 tissue types (normal or cancer).

### Architecture of MLP models

There was only one hidden layer in the MLP models with 128 units. Batch normalization and ReLU were applied immediately after the hidden layer. There were 19,241 variables in the input layer, each corresponding to one of the 19,241 genes. The output layer assigns a phenotype probability-like score for each of the 82 tissue types.

### Model training

All samples in a tissue type were randomly partitioned at 9:1 ratio, with 90% of samples used as training data and the remaining 10% as testing data. In each epoch, up-sampling was employed to avoid imbalance caused by different sample sizes between tissue types. Adam optimizer on cross entropy loss was utilized to update the weights of the neural network. After hyperparameter optimization, an initial learning rate of 0.0006 was used for CNN models, and 0.001 was used for MLP models. Batch size of 256 was used for both CNN and MLP. If there is no improvement for 5 sequential epochs, the learning rate was reduced by 0.25. L2 regularization was applied with a lambda score of 0.001. A fixed dropout of 0.25 was applied before the output layer in the MLP models, while dropout of 0.25 was applied before the global average pooling layer in the CNN models.

To optimize reproducibility in model explanation, we selected 60 well-trained models with slightly different parameters but of similar performances. In the CNN model, genes were organized in to 2-D matrix with fixed orders as input. In this study, gene orders in the CNN model were also experimented. For testing of the same gene order, we selected 5 well well-trained models with slightly different parameters but of similar performances. To study different gene orders, we selected 60 well-trained models with slightly different parameters but of similar performances.

### Estimate model performance

Five-fold cross validation was used to estimate the model performance for both MLP and CNN. Five groups of datasets were prepared, and each included a training dataset and a test dataset. Dataset preparation for each group was as following. First, we randomly split all samples in a tissue type into 5 parts. Each group used one of the 5 parts as test dataset and the remaining four parts combined into the training dataset. The same hyperparameters were used to train models based on the training dataset of each group separately. The trained models were then used to estimate the test dataset of the same group. Estimated results from five groups were combined and all metrics about performance were calculated based on the combined results.

### Model explanation

To estimate how much each gene contributes to the model prediction, we used eight different model explainers and variations, which are DeepLift, DeepLiftShap, GuidedBackprop, GuidedGradCam, GuidedGradCam++, InputXGradient, IntegratedGradients, and Saliency. All these explainers were implemented based on Captum package (https://github.com/pytorch/captum). All explanation were based on well-trained CNN and MLP models. In addition, dropout was also enabled to increase the diversity of model architecture, which helps measuring the impact of model variations and uncertainties during model explanation. As output, each explainer estimated contribution scores for each of the 19,241 genes.

### Reference preparation

In this study, we tested four kinds of references, which are named as zero, normal, universal and specific. 1) For reference zero, we assigned expression level of each gene to 0. 2) For reference norm, expression level of each gene was randomly generated from a truncated normal distribution ***N*** (0,1), and all values are restricted between 0 and 1. 3) For reference universal, expression level of each gene was randomly generated from a truncated normal distribution ***N*** (µ, σ), and all values are restricted between 0 and σ. Here, µ and σ were calculated based on expression values of a specific gene across all samples from all tissues, and σ is the standard deviation. 4) Reference specific was generated the same way as reference universal except that only samples of the same tissue types were used. In the reference testing, 2,000 different references were generated for reference normal, universal and specific separately. For zero, we just repeated the same reference 2,000 times.

### Simulation for optimal number of pseudo-samples generated for each sample

Simulation was performed on 5 different pretrained models as following, using sample X as an example. Step 1: we generated 50 pseudo-samples based on sample X by randomly adding noise to each gene’s expression level with normal distribution ***N*** (0,1). Step 2: we estimated gene contribution scores for all genes in each pseudo-sample. To estimate gene contribution scores based on *n* pseudo-samples, we randomly selected *n* replicates out of 50, and the final gene contribution score for a specific gene was calculated based on mean of *n* scores. Step 3: repeat step 1) and 2) on 5 different pretrained models respectively. Step 4: For sample X, there will be 5 replicates of gene contribution scores based on the same number of pseudo-samples. For the 5 replicates based on *n* (*n* = 1, 2, …, 100) pseudo-samples, we calculated Spearman’s Correlation coefficient on gene contribution scores from any two replicates, and this operation was carried out on all 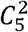 combinations. Based on the above method, we obtained the relationship between number of pseudo-samples and Correlation coefficient on any two replicates.

### Simulation for optimal repeat number on the same model

The simulation process was performed on 5 different pretrained models as following, using sample X as an example. Step 1: First, we estimated gene contribution scores for all genes in sample X 50 times respectively, and there were 50 replicates for sample X. To estimate gene contribution scores by *n* times repeats, we randomly selected *n* replicates out of 50, and the final gene contribution score for a specific gene was calculated based on mean of *n* scores. Step 2: repeat step 1) on 5 different pretrained models respectively. Step 3: For sample X, there will be 5 replicates of gene contribution scores based on the same repeat number. For the 5 replicates based on *n* (*n* = 1, 2, …, 100) times repeats, we calculated Spearman’s Correlation coefficient on gene contribution scores from any two replicates, and this operation was carried out on all 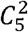 combinations. Based on the above method, we obtained the relationship between repeat number and Correlation coefficient on any two replicates.

### Simulation for optimal number of references

The simulation process was performed on 5 different pretrained models as following, using sample X as an example. Step 1: we estimated gene contribution scores for all genes in sample X with 1, 2, 3 … 100 reference samples respectively, and the reference samples were randomly selected from the 2,000 background samples pool. Step 2: repeat step 1) on 5 different pretrained models. Step 3: For sample X, there will be 5 replicates of gene contribution scores based on the same number of reference samples but different pretrained models. For the 5 replicates based on *n* (*n* = 1, 2, …, 100) reference samples, we calculated Spearman’s Correlation coefficient on gene contribution scores from any two replicates, and this operation was carried out on all 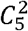 combinations. Based on the above method, we obtained the relationship between number of reference samples and Correlation coefficient on any two replicates. For each type of reference, we repeated the above simulation process individually.

### Simulation for optimal number of aggregated models

The simulation process was performed on 60 different pretrained models as following, using sample X as an example. Step 1: we estimated gene contribution scores for all genes in sample X on each pretrained models respectively. Step 2: to estimate gene contribution scores by aggregating *n* (*n* = 1, 2, …, 20) models, we randomly selected *n* replicates out of 60, and the final gene contribution score for a specific gene was calculated based on mean of *n* scores. Step 3: repeat step 2) *K* times, where 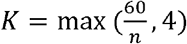. Step 4: For sample X, there will be *K* replicates of gene contribution scores based on the same number of aggregated models. For these *K* replicates based on *n* (*n* = 1, 2, …, 20) aggregated models, we calculated Spearman’s Correlation coefficient on gene contribution scores from any two replicates, and this operation was carried out on all 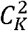 combinations. Based on the above method, we obtained the relationship between repeat number and Correlation coefficient on any two replicates.

### Gene classification

Tissue-specific expressed genes were identified by the tool TissueEnrich with group “Tissue-Enhanced” [45]. In each tissue, median expression level of each gene was calculated across all samples, and housekeeping genes are defined as genes with TPM ≥1 and less than 2-fold change on median expression level among all tissue types [46].

### Gene Ontology enrichment analysis

Genes of interest were extracted and imported into the Gene Ontology online tool for Gene Ontology enrichment analysis with the options “molecular function” or “biological process” and “Homo sapiens” checked [47, 48].

### Annotation of transcription factor (TF) and TF cofactors

All TFs and TF cofactors were downloaded from animalTFDB [49]. In total, there were 1,666 TFs and 1,026 TF cofactors.

### Differentially expressed genes between normal and cancer

Mann–Whitney U test (two-sided) was used to compare gene expression between normal and cancer tissues. Differentially expressed genes should satisfy the following criteria: FDR ≤ 0.001 and fold change ≥ 3.

## Legends

**Figure S1 Performance of model neural network interpretability.**

**A**. Spearman’s correlation on gene contribution scores and **B**. overlap in the top 100 contributing genes in lung (upper panel), ovary (middle panel), and pancreas (lower panel) among replicates from the same pretrained model, different pretrained models with the same gene order, and different pretrained models with different gene orders based on CNN. **C**. Spearman’s correlation on gene contribution scores and **D**. overlap in the top 100 contributing genes in lung (upper panel), ovary (middle panel), and pancreas (lower panel) among replicates from the same pretrained model and different pretrained models based on MLP.

**Figure S2 Performance of repeat with adding noise**.

Spearman’s correlation on gene contribution scores in lung (left panel), ovary (middle panel), and pancreas (right panel) among replicates from **A**. different pretrained models with different gene orders on CNN and from **B**. different pretrained models on MLP.

**Figure S3 Performance of simple repeat**.

Spearman’s correlation on gene contribution scores in lung (left panel), ovary (middle panel), and pancreas (right panel) among replicates from **A**. different pretrained models with different gene orders on CNN and from **B**. different pretrained models on MLP.

**Figure S4 Performance of reference type**.

Spearman’s correlation on gene contribution scores among replicates from **A**. different pretrained models with different gene orders on CNN and from **B**. different pretrained models on MLP.

**Figure S5 Performance of different reference types with simple repeat**.

Spearman’s correlation on gene contribution scores among replicates from different pretrained models with different gene orders on CNN.

**Figure S6 Performance of model aggregation**.

Spearman’s correlation on gene contribution scores in lung (left panel), ovary (middle panel), and pancreas (right panel) among replicates from **A**. different pretrained models with different gene orders on CNN and from **B**. different pretrained models on MLP.

**Figure S7 Optimization on different model explainers.**

**A**. Spearman’s correlation on gene contribution scores and **B**. overlap in the top 100 contributing genes in lung (upper panel), ovary (middle panel), and pancreas (lower panel) among replicates from different pretrained models with different gene orders on CNN. The analyses were carried out with different optimization strategies: without optimization, with optimized conditions for each explainer but without model aggregation, and with optimized conditions for each explainer and with model aggregation. **C**. and **D**. similar analysis as A and B but based on MLP.

**Figure S8 Overlap between the top contributing genes with and without optimization**.

Comparing top 100 contributing genes with optimized method with the top 100 contributing genes in individual replicates without optimization (dark blue) and top 100 contributing genes shared in 2 or more replicates without optimization (yellow).

**Figure S9 Spearman’s correlation between gene contribution scores and gene expression level**.

Left panels are analyses based on CNN with and without model aggregation, while right panels are based on MLP.

**Figure S10 Overlap between the top 100 contributing genes and top 100 expressed genes**.

Left panels are analyses based on CNN with and without model aggregation, while right panels are based on MLP.

**Figure S11 Overlap between the top 100 contributing genes and tissue-specifically (TS) expressed genes.**

**A**. Left panels are analyses based on CNN with and without model aggregation, while right panels are based on MLP. **B**. Number of TS genes in liver, lung, ovary and pancreas.

**Figure S12 Overlap between the top 100 contributing genes and housekeeping (HK) genes**.

Left panels are analyses based on CNN with and without model aggregation, while right panels are based on MLP.

**Figure S13 Overlap between the top 100 contributing genes and transcription factors (TFs)**.

Left panels are analyses based on CNN with and without model aggregation, while right panels are based on MLP.

**Figure S14 Overlap between the top 100 contributing genes and transcription factor (TF) cofactors**.

Left panels are analyses based on CNN with and without model aggregation, while right panels are based on MLP.

**Figure S15 Expression level of the top 100 contributing genes between normal and cancer tissues**.

**Figure S16 Heatmap of 45 tumor oncogenes and suppressor genes identified as shared top contributing genes across 33 cancer types**.

**Table S1 Information for different model explainers**.

**Table S2 Information for 82 different tissues and cell types used in model training**.

**Table S3 Overlaps in the top 100 contributing genes across different explainers**.

**Table S4 Overlaps in the top 100 contributing genes across tissues in different explainers**.

**Table S5 GO enrichment on molecular function on the top 100 genes in different explainers**.

