## Supplementary Figures for "Assessment and Optimization of the Interpretability of Machine Learning Models Applied to Transcriptomic Data"

### Figure S1

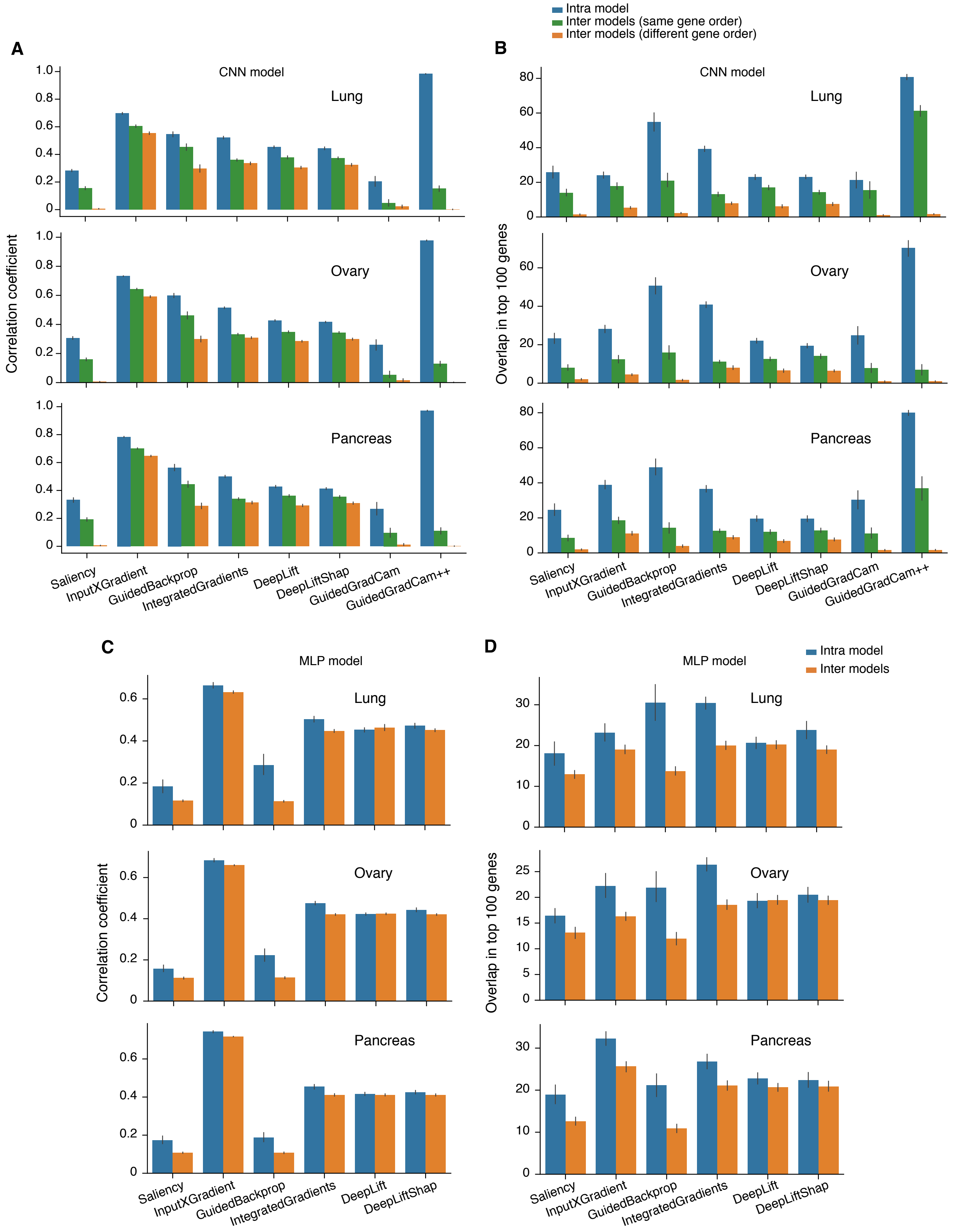

Intra model
Inter models (same gene order)
Inter models (different gene order)
A
B
CNN model
Lung
Ovary
Pancreas
Correlation coefficient
Overlap in top 100 genes
Saliency
InputXGradient
GuidedBackprop
IntegratedGradients
DeepLift
DeepLiftShap
GuidedGradCam
GuidedGradCam++
C
D
MLP model
Intra model
Inter models

### Figure S2

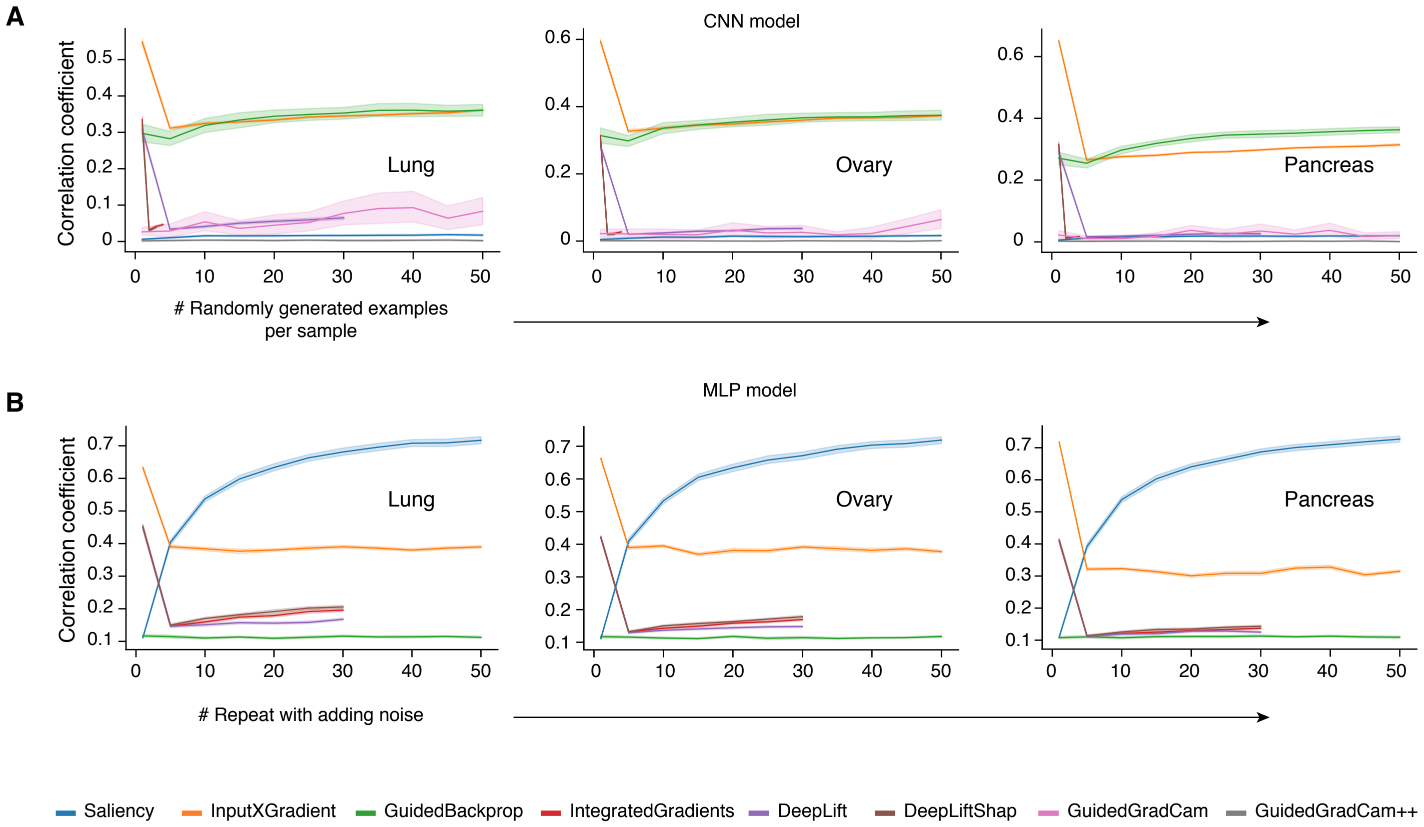
A
CNN model
Correlation coefficient
Lung
Ovary
Pancreas
### Randomly generated examples per sample
B
MLP model
Correlation coefficient
Lung
Ovary
Pancreas
### Repeat with adding noise
Saliency
InputXGradient
GuidedBackprop
IntegratedGradients
DeepLift
DeepLiftShap
GuidedGradCam
GuidedGradCam++

### Figure S3

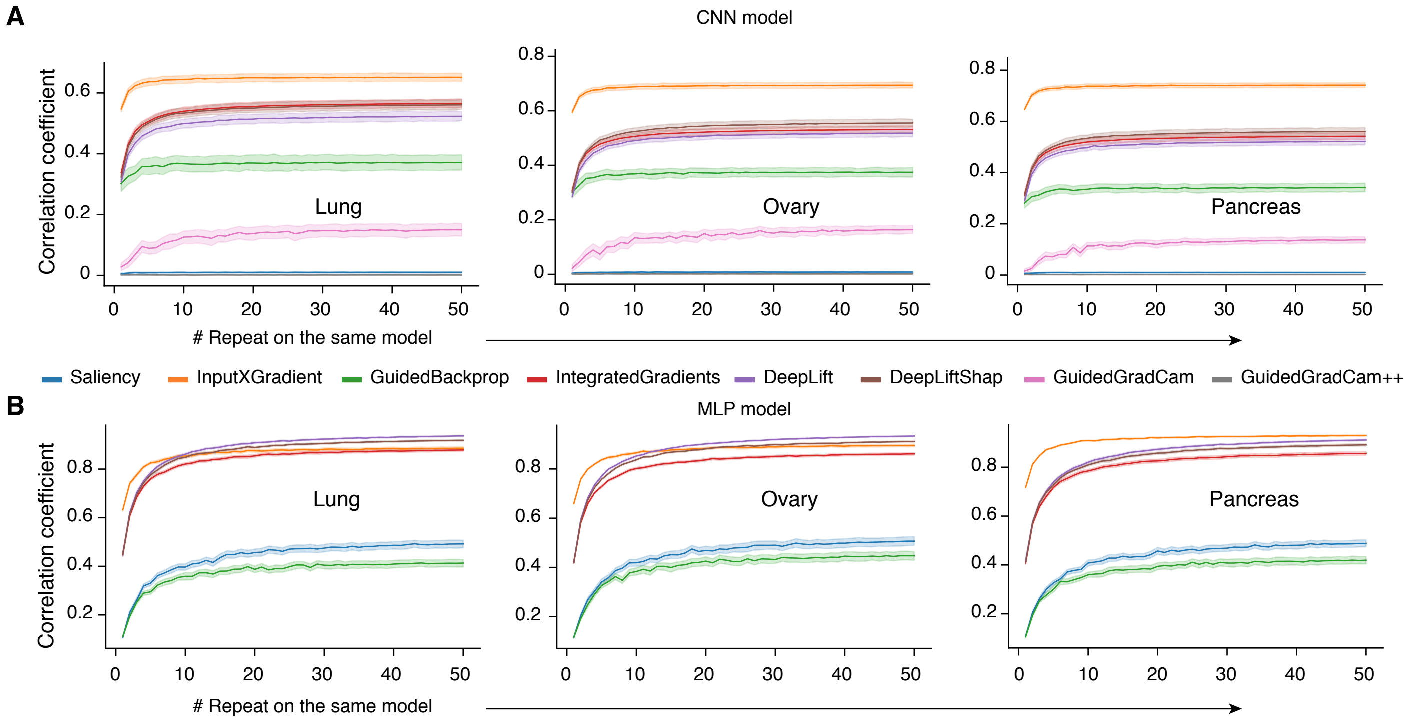

A
CNN model
Correlation coefficient
Lung
Ovary
Pancreas
### Repeat on the same model
Saliency
InputXGradient
GuidedBackprop
IntegratedGradients
DeepLift
DeepLiftShap
GuidedGradCam
GuidedGradCam++
B
MLP model
Correlation coefficient
Lung
Ovary
Pancreas
### Repeat on the same model

### Figure S4

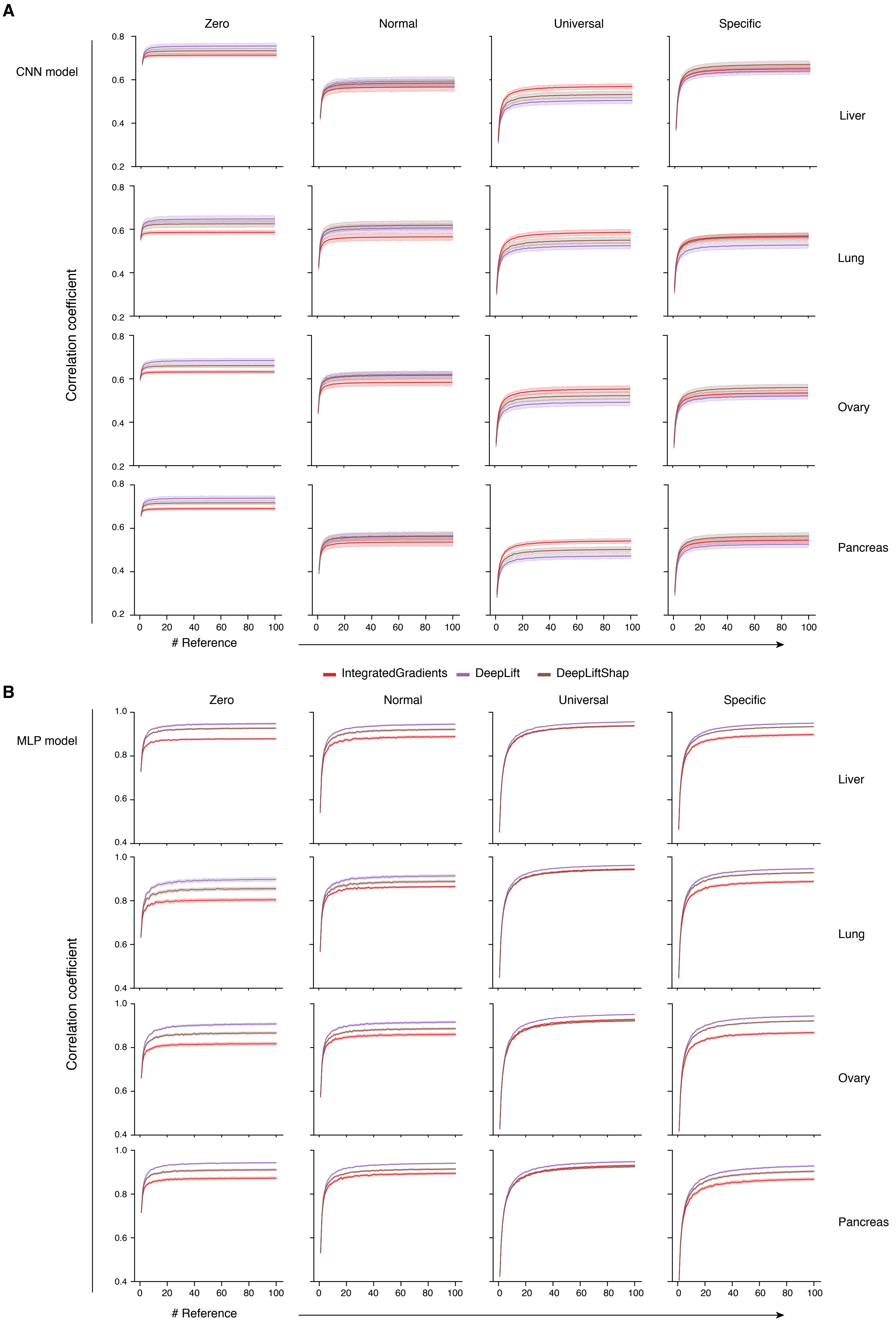

A
Zero
Normal
Universal
Specific
CNN model
Correlation coefficient
Liver
Lung
Ovary
Pancreas
### Reference
IntegratedGradients
DeepLift
DeepLiftShap
B
MLP model

### Figure S5

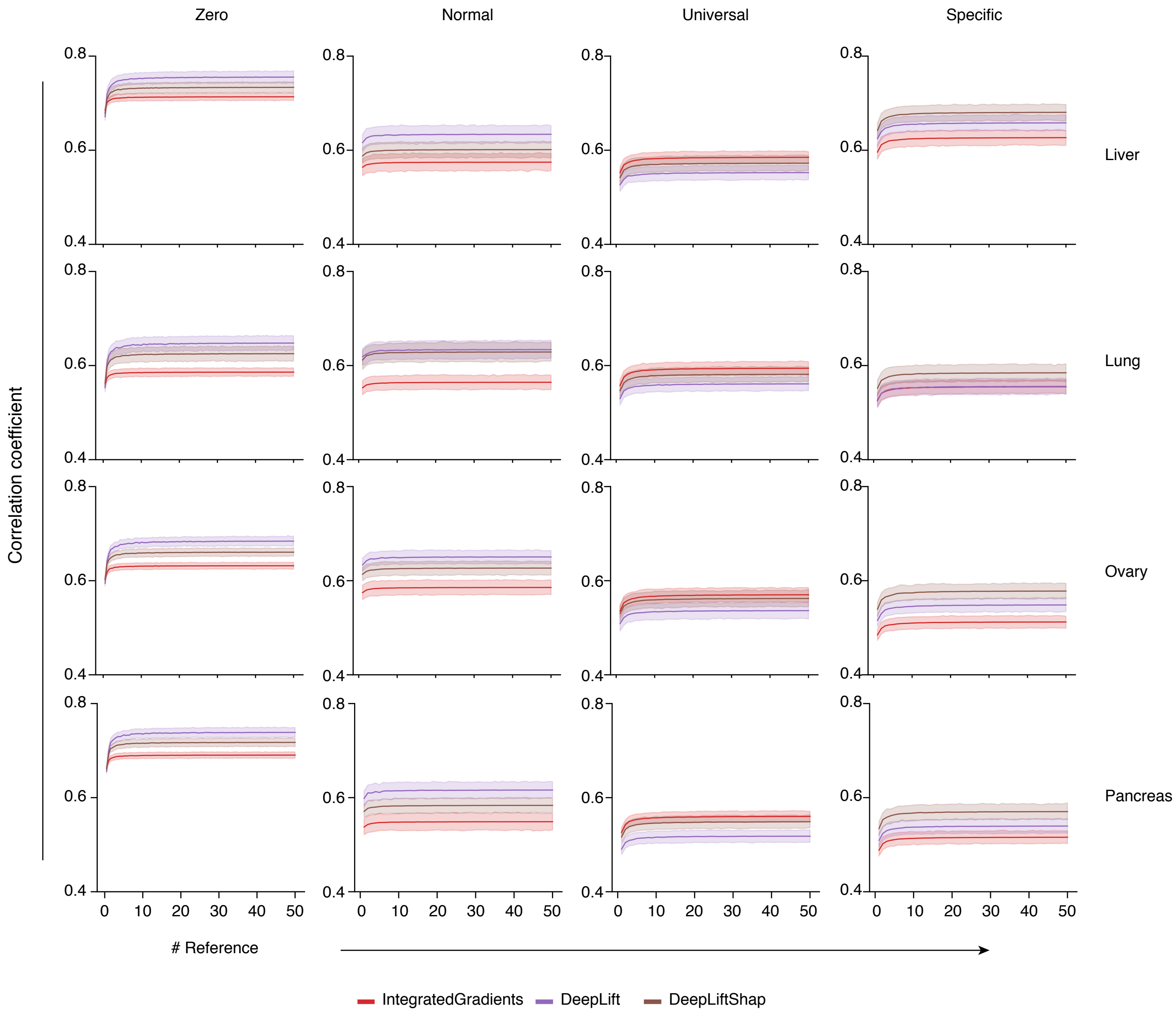
Zero
Normal
Universal
Specific
Liver
Lung
Ovary
Pancreas
Correlation coefficient
0.8
0.6
0.4
0
10
20
30
40
50
### Reference
IntegratedGradients
DeepLift
DeepLiftShap

### Figure S6

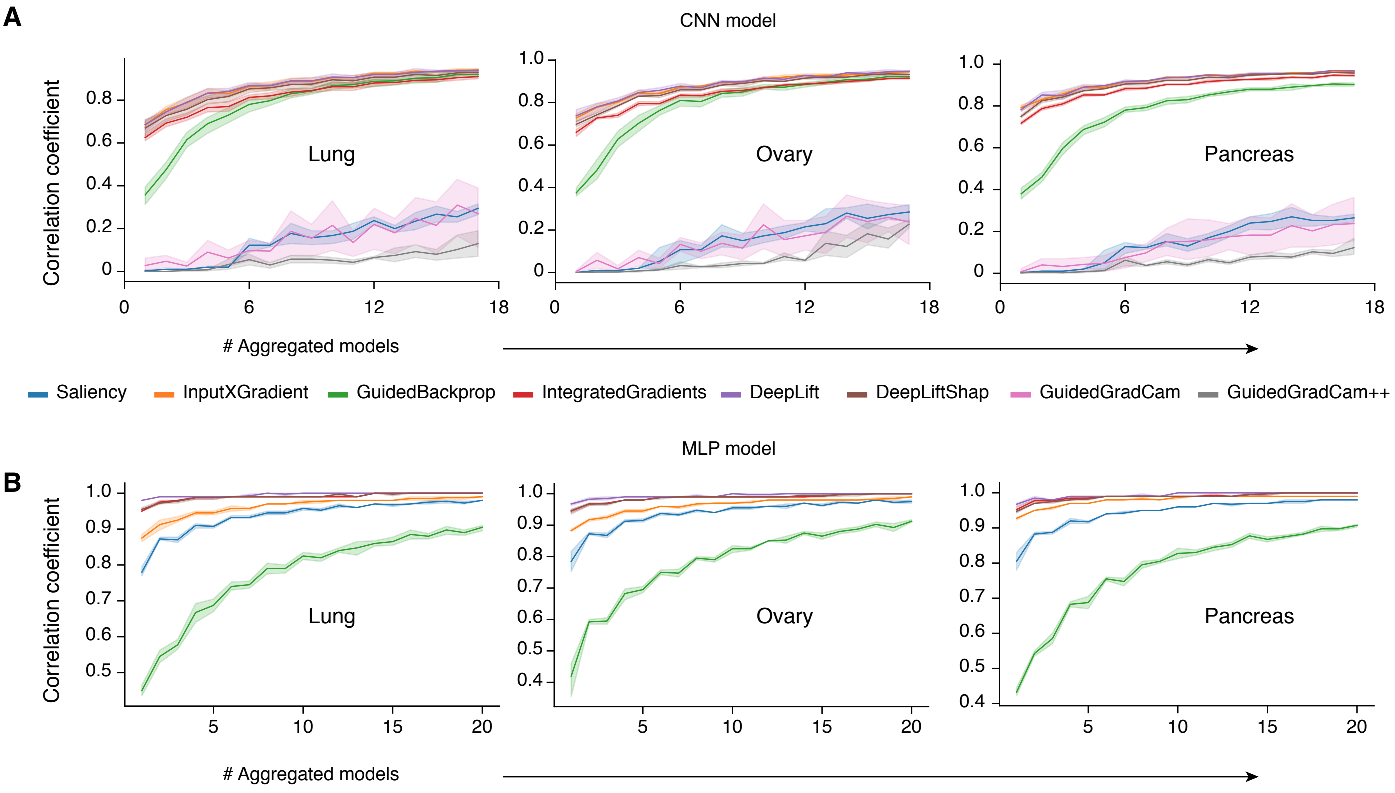
A
CNN model
Correlation coefficient
Lung
Ovary
Pancreas
### Aggregated models
Saliency
InputXGradient
GuidedBackprop
IntegratedGradients
DeepLift
DeepLiftShap
GuidedGradCam
GuidedGradCam++
MLP model
B
Correlation coefficient
Lung
Ovary
Pancreas
### Aggregated models

### Figure S7

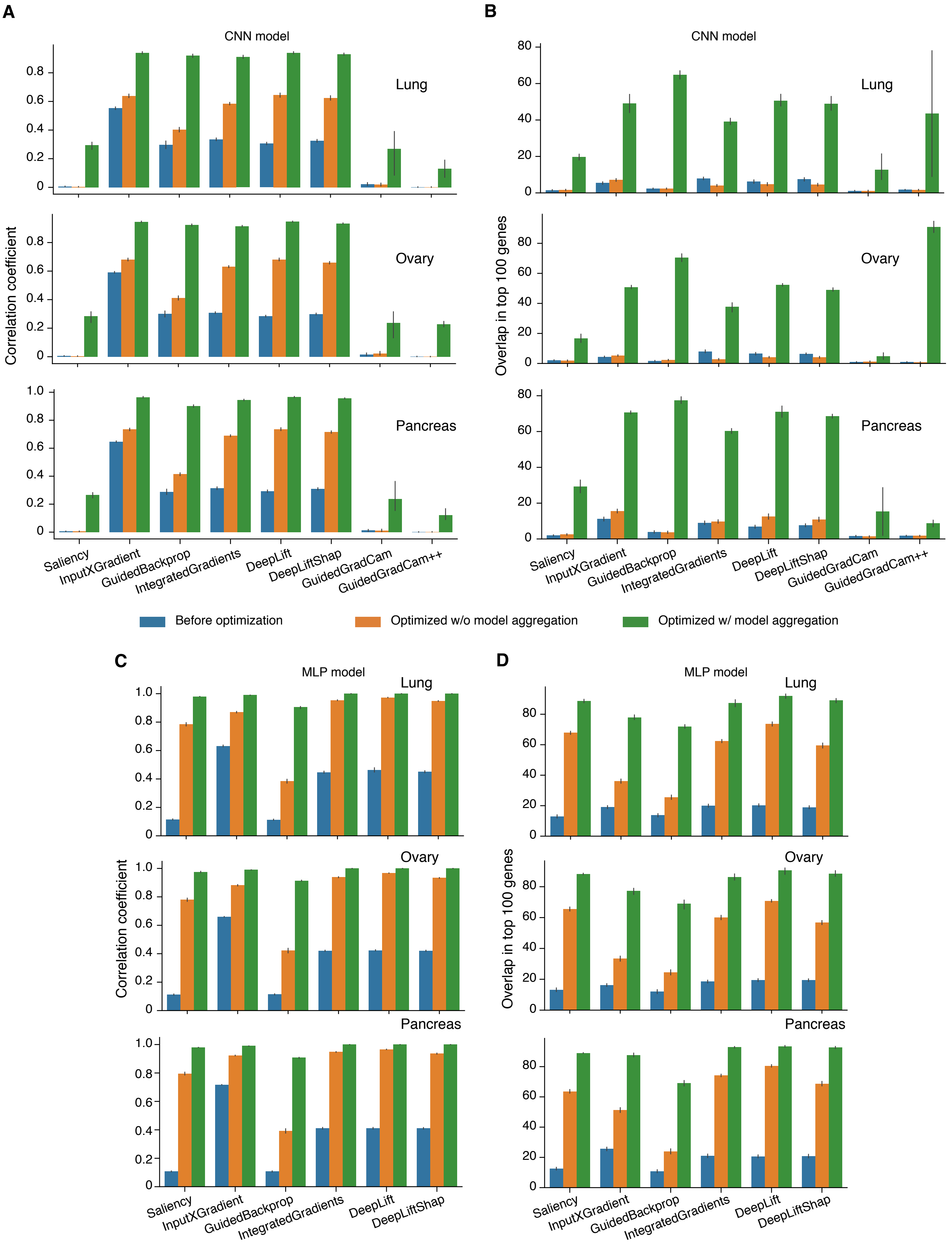

A
B
CNN model
Lung
Ovary
Pancreas
Correlation coefficient
Overlap in top 100 genes
Saliency
InputXGradient
GuidedBackprop
IntegratedGradients
DeepLift
DeepLiftShap
GuidedGradCam
GuidedGradCam++
Before optimization
Optimized w/o model aggregation
Optimized w/ model aggregation
C
D
MLP model

### Figure S8

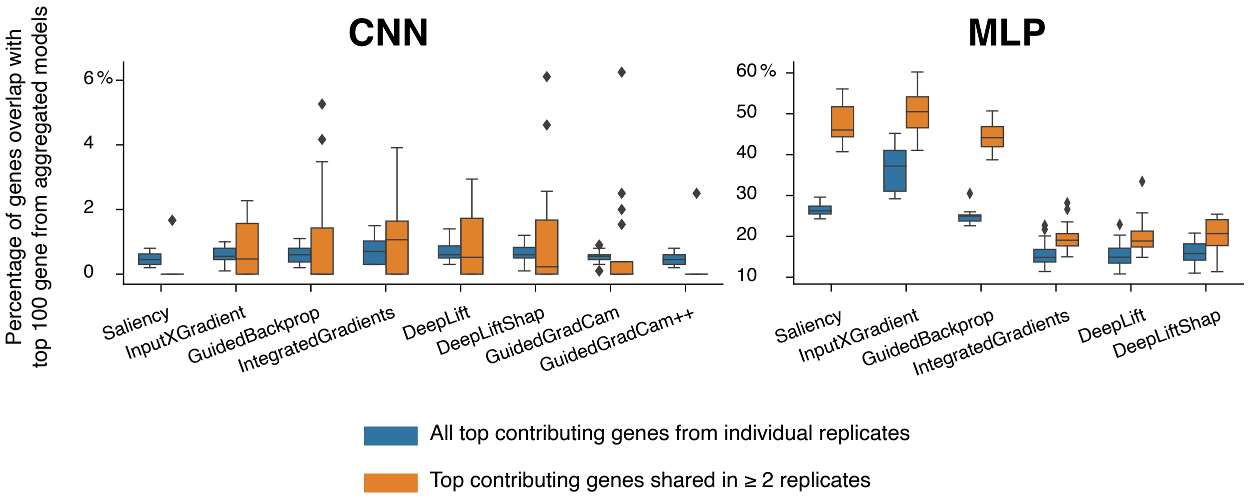
CNN
MLP
Percentage of genes overlap with top 100 gene from aggregated models
6 %
4
2
0
60 %
50
40
30
20
10
Saliency
InputXGradient
GuidedBackprop
IntegratedGradients
DeepLift
DeepLiftShap
GuidedGradCam
GuidedGradCam++
All top contributing genes from individual replicates
Top contributing genes shared in ≥ 2 replicates

### Figure S9

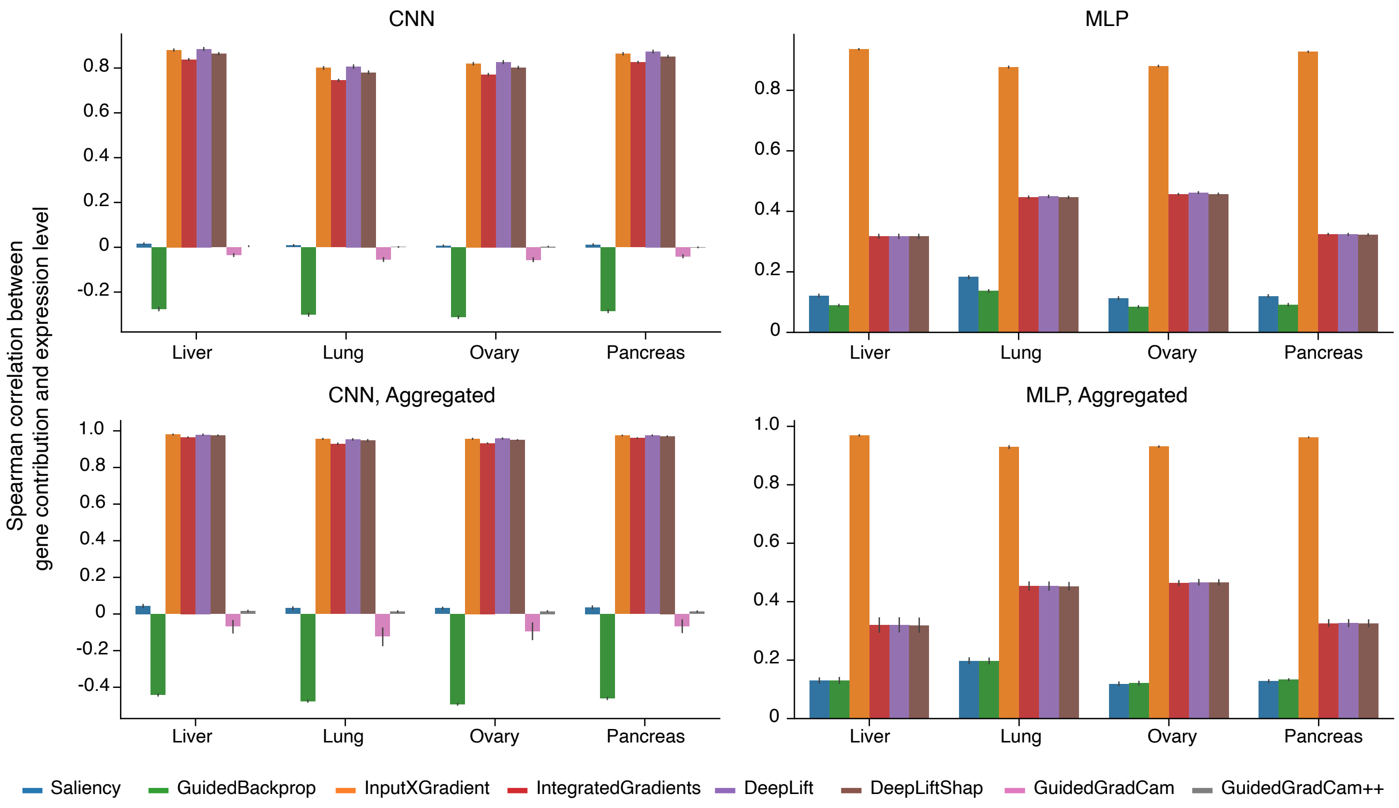
CNN
MLP
CNN, Aggregated
MLP, Aggregated
Spearman correlation between gene contribution and expression level
Liver
Lung
Ovary
Pancreas
Saliency
GuidedBackprop
InputXGradient
IntegratedGradients
DeepLift
DeepLiftShap
GuidedGradCam
GuidedGradCam++

### Figure S10

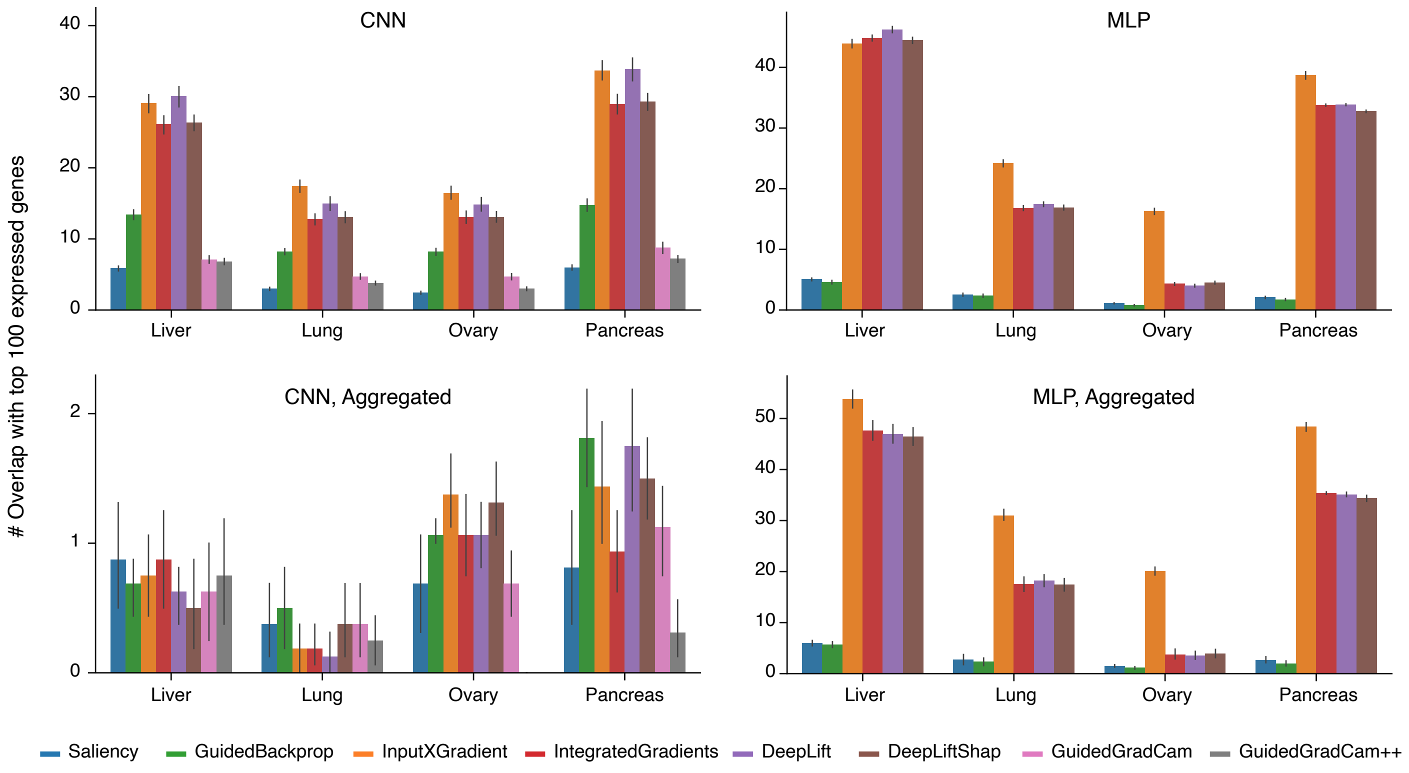

CNN
MLP
CNN, Aggregated
MLP, Aggregated
### Overlap with top 100 expressed genes
Liver
Lung
Ovary
Pancreas
Saliency
GuidedBackprop
InputXGradient
IntegratedGradients
DeepLift
DeepLiftShap
GuidedGradCam
GuidedGradCam++

### Figure S11

A

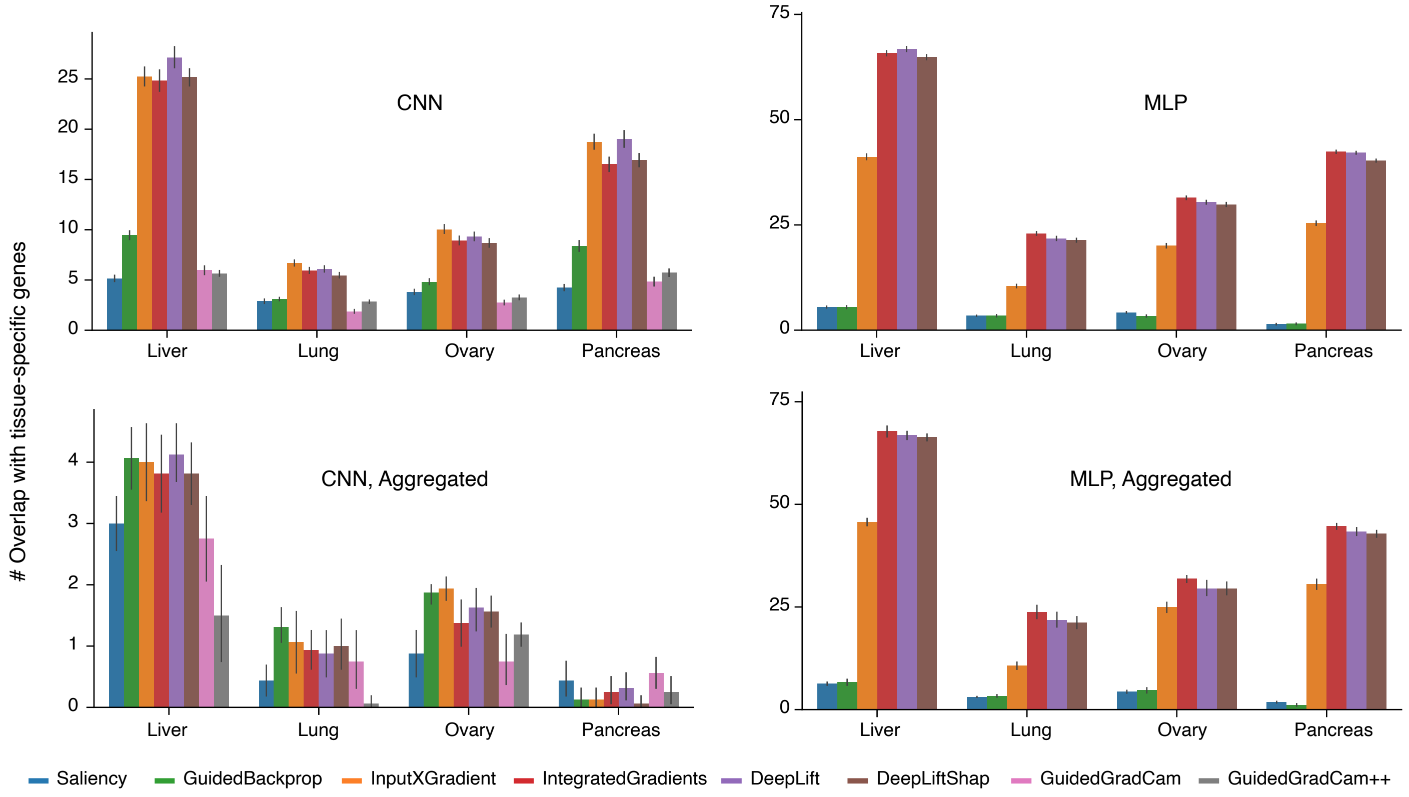


B

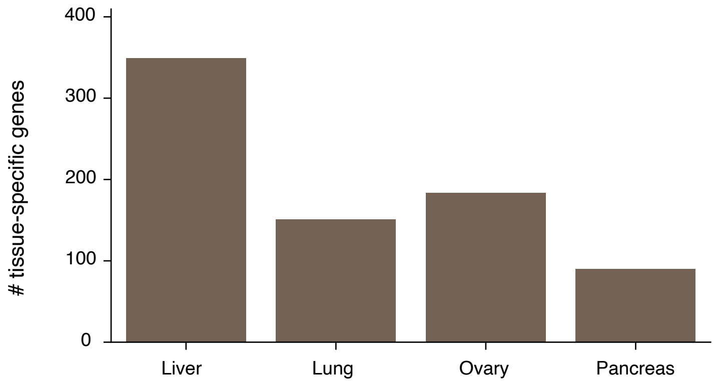

### Figure S12

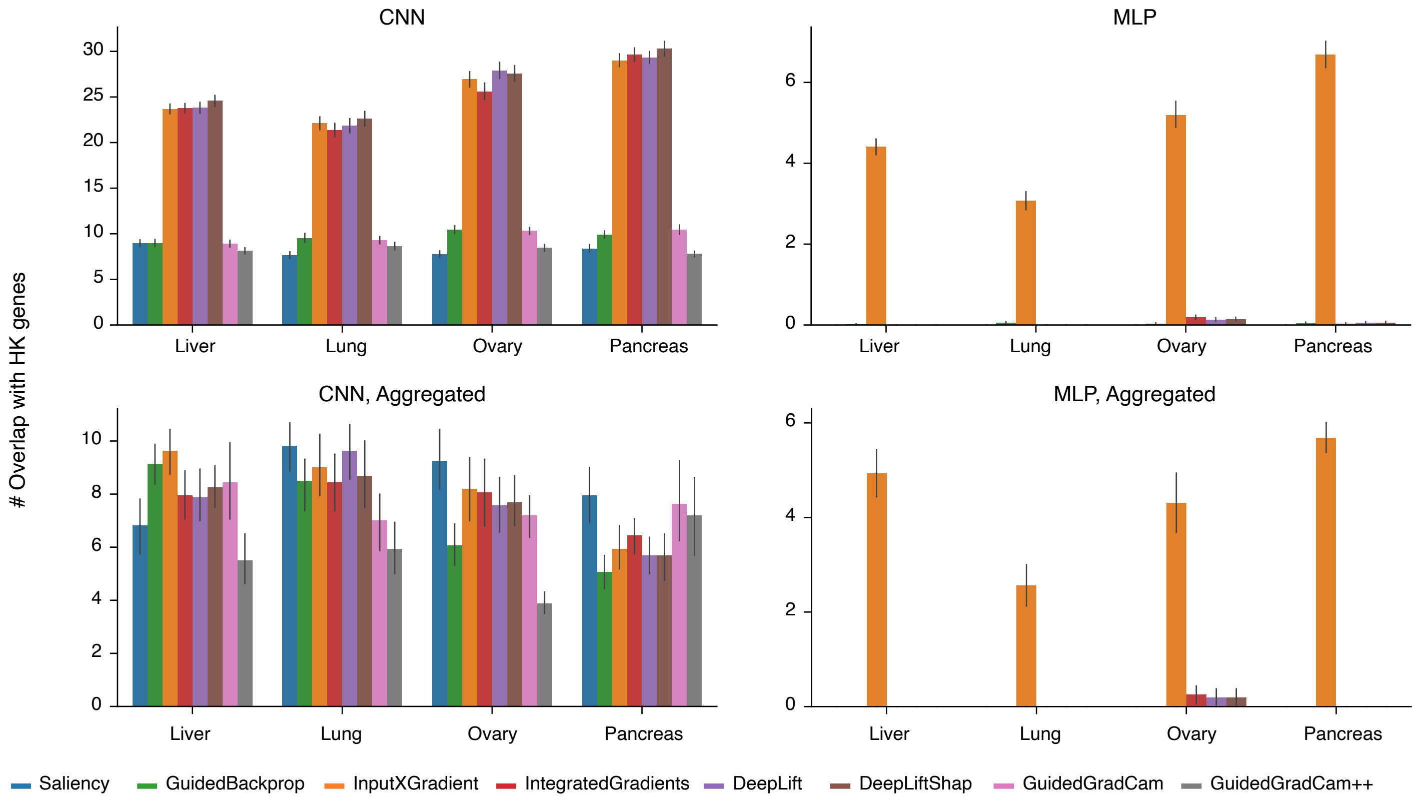

CNN
MLP
CNN, Aggregated
MLP, Aggregated
### Overlap with HK genes
Liver
Lung
Ovary
Pancreas
Saliency
GuidedBackprop
InputXGradient
IntegratedGradients
DeepLift
DeepLiftShap
GuidedGradCam
GuidedGradCam++

### Figure S13

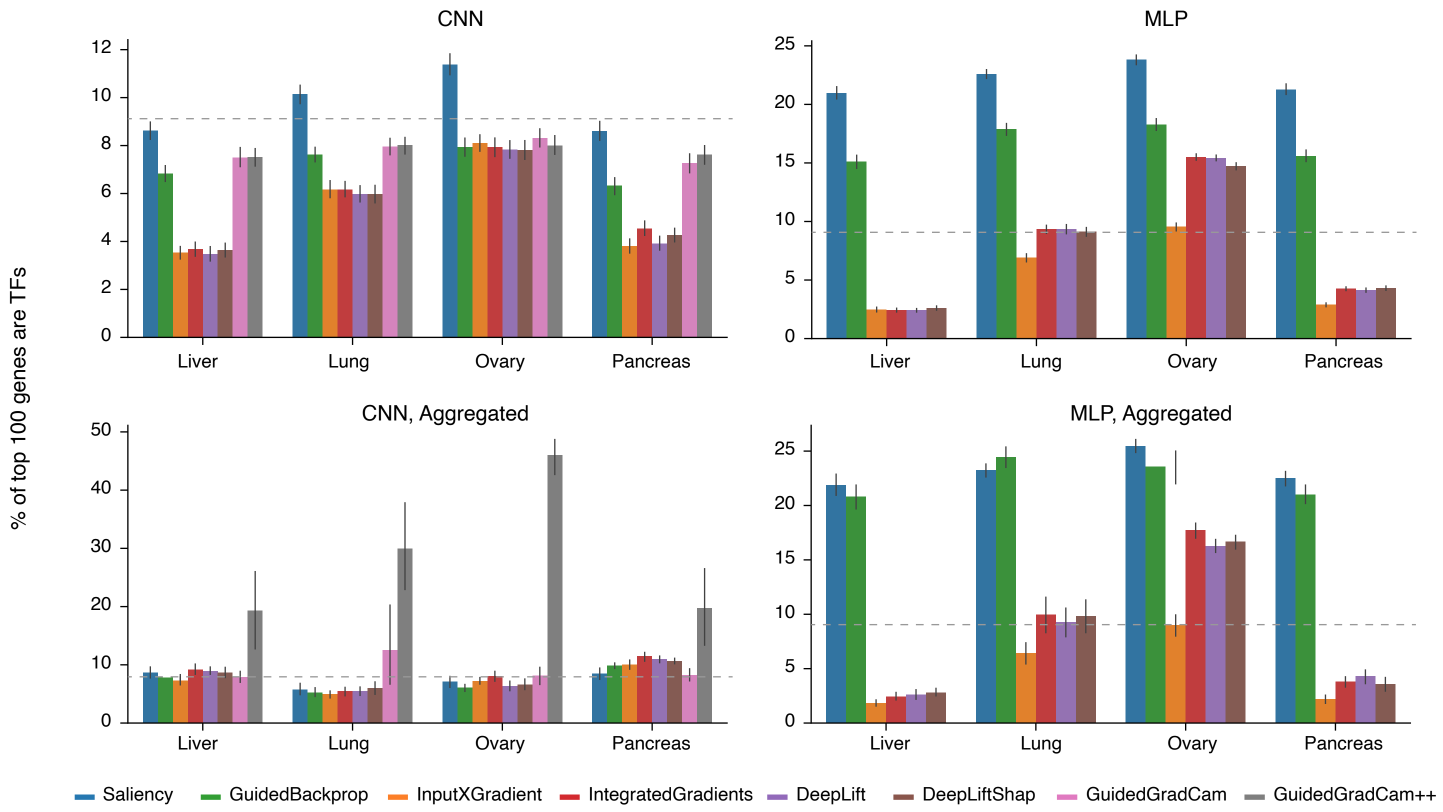
CNN
MLP
CNN, Aggregated
MLP, Aggregated
% of top 100 genes are TFs
Liver
Lung
Ovary
Pancreas
Saliency
GuidedBackprop
InputXGradient
IntegratedGradients
DeepLift
DeepLiftShap
GuidedGradCam
GuidedGradCam++

### Figure S14

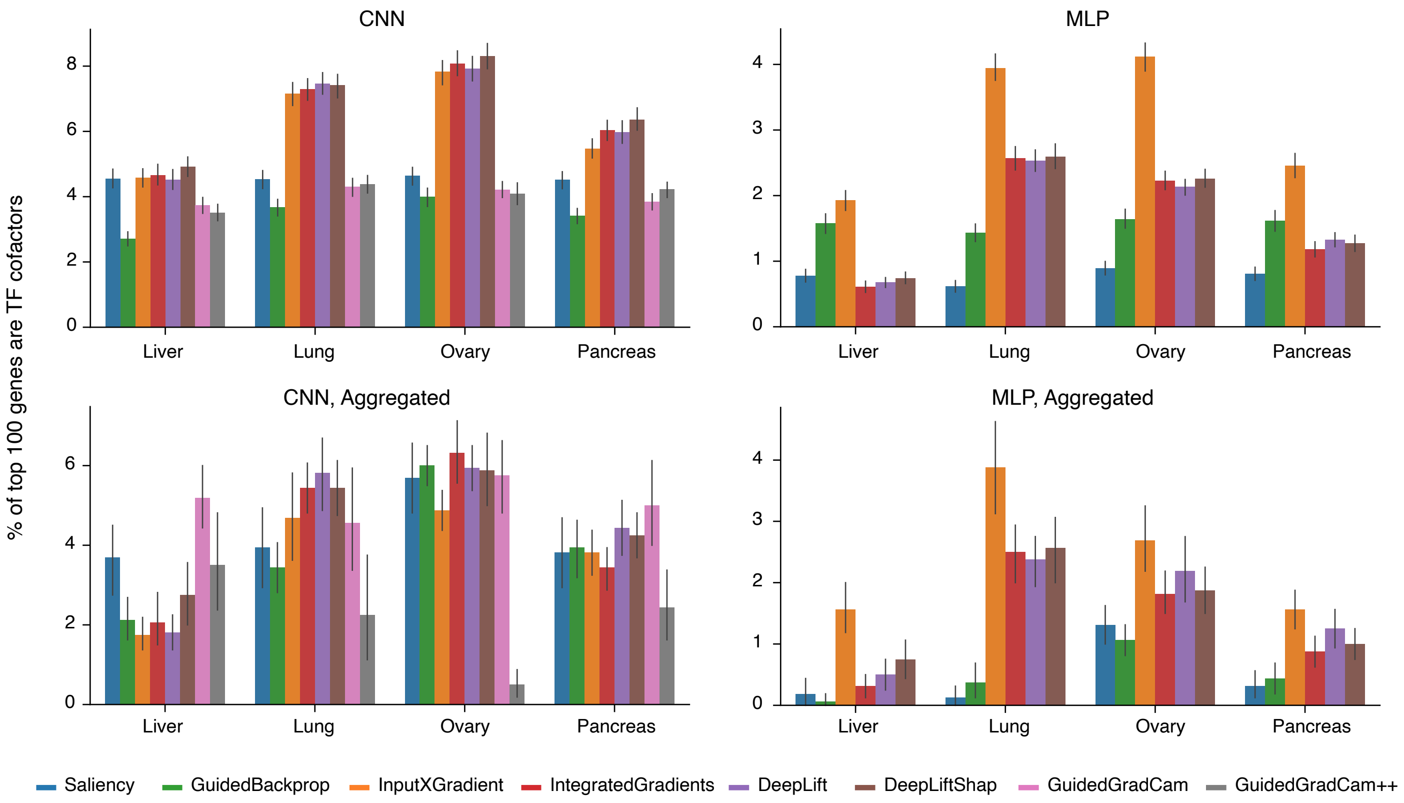

CNN
MLP
CNN, Aggregated
MLP, Aggregated
% of top 100 genes are TF cofactors
Liver
Lung
Ovary
Pancreas
Saliency
GuidedBackprop
InputXGradient
IntegratedGradients
DeepLift
DeepLiftShap
GuidedGradCam
GuidedGradCam++

### Figure S15

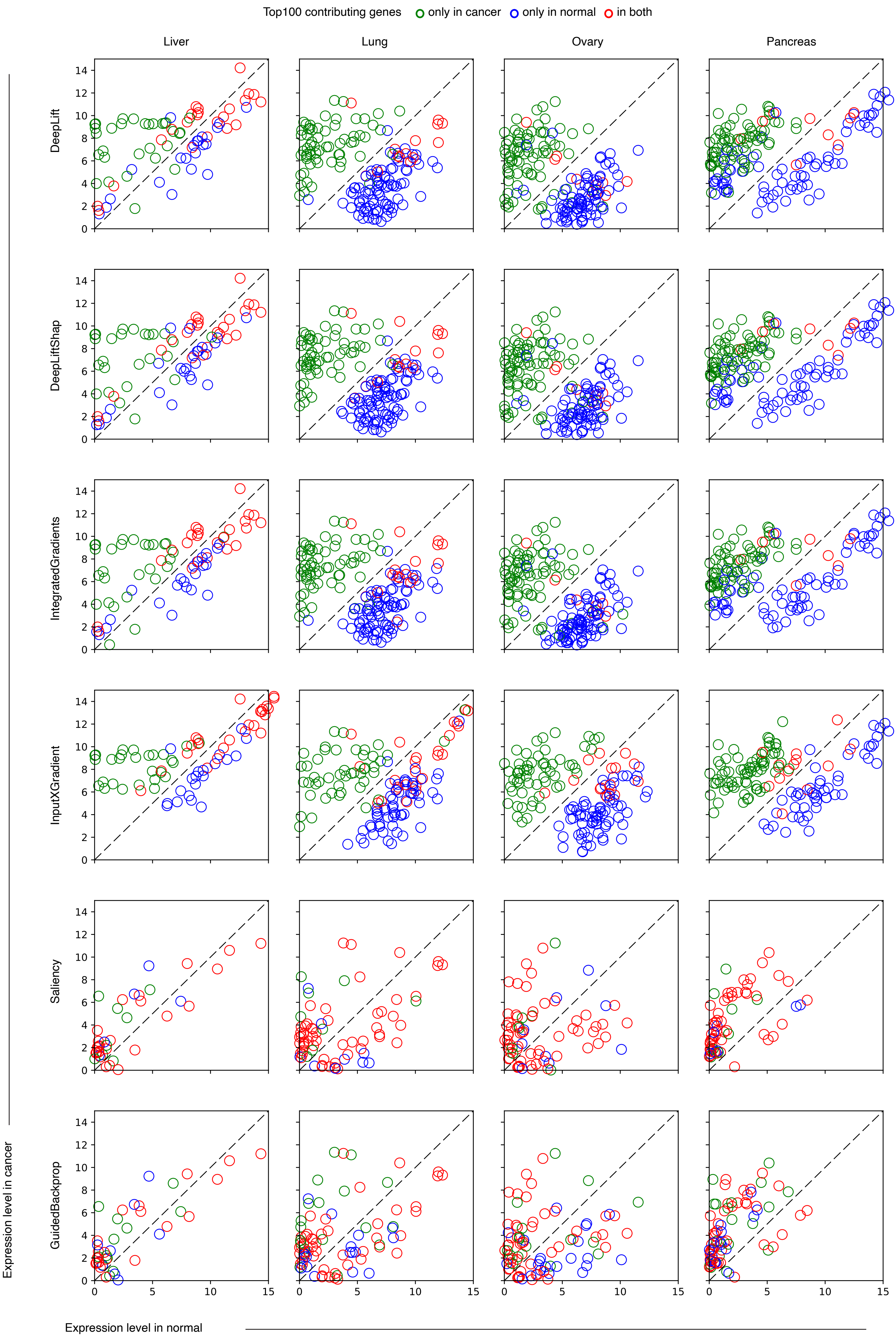
Top100 contributing genes
only in cancer
only in normal
in both
Liver
Lung
Ovary
Pancreas
DeepLift
DeepLiftShap
IntegratedGradients
InputXGradient
Saliency
GuidedBackprop
Expression level in cancer
Expression level in normal
0
2
4
6
8
10
12
14
5
10
15

### Figure S16

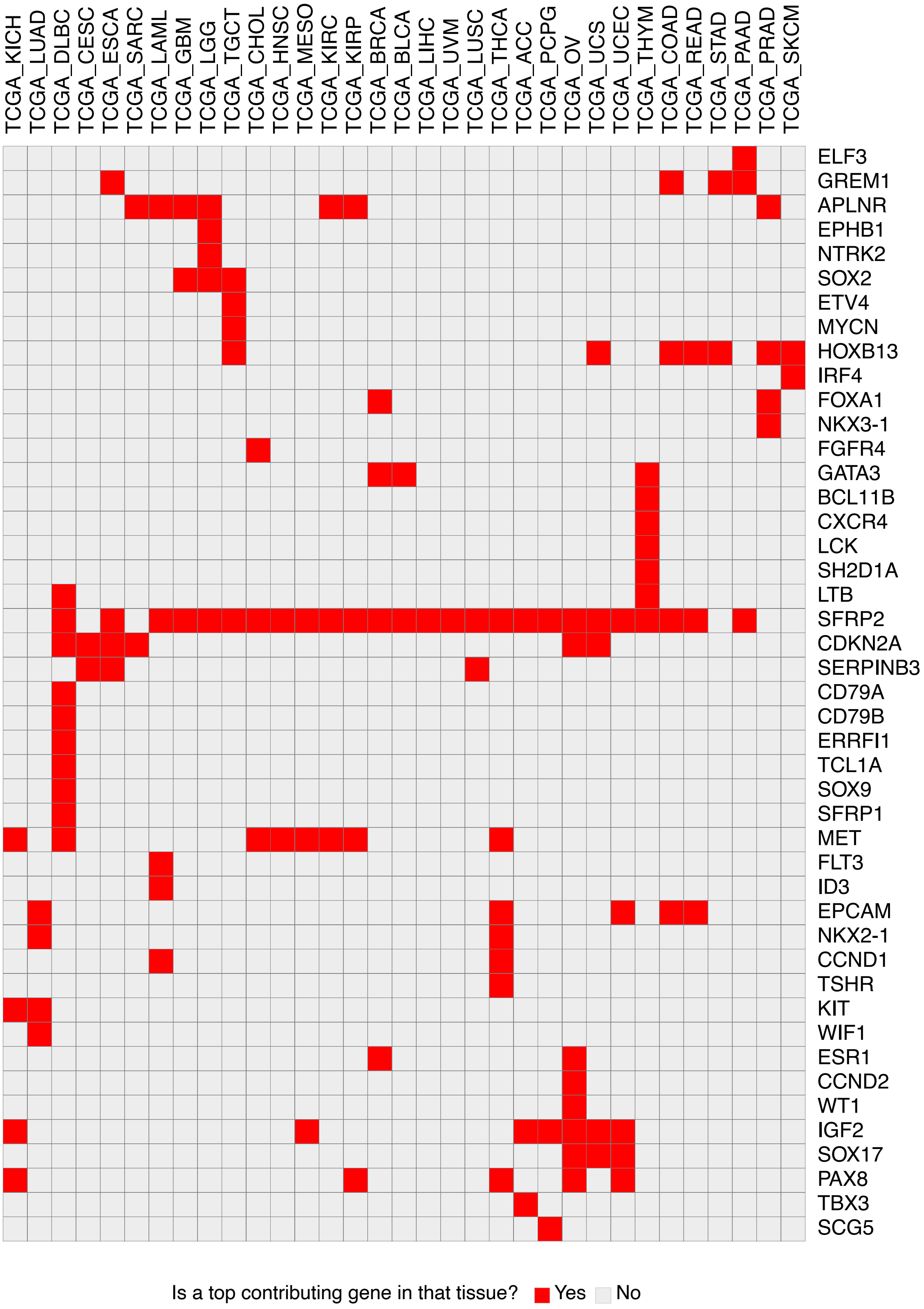
TCGA_KICH
TCGA_LUAD
TCGA_DLBC
TCGA_CESC
TCGA_ESCA
TCGA_SARC
TCGA_LAML
TCGA_GBM
TCGA_LGG
TCGA_TGCT
TCGA_CHOL
TCGA_HNSC
TCGA_MESO
TCGA_KIRC
TCGA_KIRP
TCGA_BRCA
TCGA_BLCA
TCGA_LIHC
TCGA_UVM
TCGA_LUSC
TCGA_THCA
TCGA_ACC
TCGA_PCPG
TCGA_OV
TCGA_UCS
TCGA_UCEC
TCGA_THYM
TCGA_COAD
TCGA_READ
TCGA_STAD
TCGA_PAAD
TCGA_PRAD
TCGA_SKCM
ELF3
GREM1
APLNR
EPHB1
NTRK2
SOX2
ETV4
MYCN
HOXB13
IRF4
FOXA1
NKX3-1
FGFR4
GATA3
BCL11B
CXCR4
LCK
SH2D1A
LTB
SFRP2
CDKN2A
SERPINB3
CD79A
CD79B
ERRFI1
TCL1A
SOX9
SFRP1
MET
FLT3
ID3
EPCAM
NKX2-1
CCND1
TSHR
KIT
WIF1
ESR1
CCND2
WT1
IGF2
SOX17
PAX8
TBX3
SCG5
Is a top contributing gene in that tissue?
Yes
No
